# DiffHiChIP: Identifying differential chromatin contacts from HiChIP data

**DOI:** 10.1101/2025.01.14.633096

**Authors:** Sourya Bhattacharyya, Daniela Salgado Figueroa, Katia Georgopoulos, Ferhat Ay

## Abstract

High-resolution conformation capture assays such as HiChIP are commonly used for profiling chromatin loops among *cis*-regulatory elements including enhancers and promoters. Detection of differential loops between two conditions (e.g., same cell type different genotypes or before/after perturbations) help contextualize condition-specific activities of genes in connection with such *cis*-regulatory elements. Existing differential loop callers for HiChIP data employ count-based models that are designed with gene expression data in mind and, hence, do not account for the distance decay of contact counts from HiChIP data. These approaches are not ideal for detection of differential long-range (>400Kb) loops, a limitation that persists even with the use of implicit or explicit corrections for this distance effect. We have implemented DiffHiChIP, the first comprehensive framework to call differential loops from HiChIP and similar 3C protocols. DiffHiChIP supports both DESeq2 and edgeR using either complete contact map or a subset of contacts (filtered) for background estimation, incorporates edgeR with generalized linear model (GLM) using either quasi-likelihood F-test or likelihood ratio test, and implements independent hypothesis weighting (IHW) as well as a distance stratification technique for modeling distance decay of contacts in estimating their statistical significance. Our results on 5 different datasets, each with two conditions or cell types, suggest that edgeR GLM-based models with IHW correction capture differential interactions, including long-range, that are supported by published Hi-C data and reference studies. Given the increasing trend of generating and utilizing HiChIP data for modeling chromatin regulation, DiffHiChIP promises to have broad impact and utility in this field.

## INTRODUCTION

Chromosome conformation capture (3C) technologies like Hi-C^1,2^ and its variants such as Promoter Capture Hi-C (PCHi-C)^3,4^, ChIA-PET^5^, Micro-C^6^ and others produce high-resolution 3D chromatin interaction maps of the genome^7^. One variant, commonly referred to as HiChIP (Hi-C coupled with chromatin immunoprecipitation or proximity ligation assisted ChIP-seq)^8–10^ profiles protein- or histone modification-centric (e.g., CTCF or H3K27ac) interactions requiring much lower sequencing depth than Hi-C (∼300M HiChIP reads instead of >1B Hi-C reads for 5kb resolution). This allows more scalable and cost-effective studies of genome-wide chromatin interactions or loops (we reserve the term loop for interactions that meet certain significance criteria) across regulatory and/or structural elements from multiple different cell types and conditions^11–15^. The increasing availability of HiChIP datasets also prompt the need to systematically identify their similarities and differences across conditions. Although various methods have assessed the reproducibility^16–18^ and differential loops^19–22^ from Hi-C and PCHi-C contact maps, similar studies for HiChIP data are limited^23–26^. This gap is partly due to the challenges of modeling the non-uniform coverage of HiChIP contact counts according to the underlying ChIP-seq (1D) signal and the genomic distance between interacting loci. Existing differential HiChIP callers such as diffloop^25^, HiCDC+^24^ and FitHiChIP^23^ have employed RNA-seq count based techniques DESeq2^27^ and edgeR^28^. While DESeq2 employs negative binomial (NB) distribution on the input count matrix and generalized linear model (GLM) based regression to estimate the gene-wise dispersions, edgeR exactTest uses the percentiles (or rank) of gene expression to compute gene-wise dispersions and estimate gene-specific p-values using the quantile normalized RNA-seq counts^29^. However, HiChIP contacts exhibit much higher dispersion than 1D RNA-seq and ChIP-seq assays due to inherent 3C-based biases (e.g., position of restriction fragments, distance effect) as well as underlying ChIP-seq signals^30,31^. We reasoned that GLM-based regression in edgeR employing both *gene-wise* and *common* (or *trended*) dispersion together with likelihood ratio test (LRT)^29^ or quasi-likelihood-F test (QLFTest)^32^ would likely fit the HiChIP contacts better than the exactTest setting, akin to their application in estimating differential abundance of single cell clusters^33,34^. Additionally, neither DESeq2 nor edgeR model the exponential distance decay of chromatin contacts^35,36^, thus mostly ignoring the longer-range (>400Kb) differential loops. To model such distance decay in estimating the p-values of chromatin contacts, previous studies have either employed *Independent Hypothesis Weighting* (IHW)^37^ on DESeq2 p-values^38^ or applied *distance stratification* to first distribute the chromatin contacts into different bins subject to their genomic distances, and then estimate statistical significance separately for individual bins^24,39^. Relative utilities of these distance decay modeling techniques remain to be systematically benchmarked with comprehensive datasets and metrics that utilize orthogonal data to assess their accuracy.

Here we present DiffHiChIP, a comprehensive framework to identify differential HiChIP loops by integrating various count-based approaches (DESeq2 or edgeR), supporting multiple dispersion estimation techniques (exactTest or GLM), employing different statistical tests (LRT, QLFTest), and modeling distance decay in multiple ways (e.g., IHW, distance stratification). We provide a comprehensive assessment of each of these aspects of differential HiChIP analysis using 5 different HiChIP datasets spanning perturbations of regulators of chromatin looping, cytokine stimulation and different cell types. We also utilize matched data from Hi-C, ChIP-seq and RNA-seq experiments in these conditions as well as gene/loci highlighted in these studies for evaluating differential loop calls. Our results suggest that: **1)** IHW correction of p-values generally performs better in capturing longer-range differential HiChIP loops compared to BH correction or distance stratification, **2)** generalized linear model (GLM)-based statistical tests in edgeR exhibit higher sensitivity of differential loop calling than DESeq2 and edgeR exactTest models, particularly for datasets with lower number of replicates (n=2), **3)** For datasets with higher number of replicates per condition, DESeq2 reports considerably higher number of differential loops but with lower specificity. Although the results vary substantially across different HiChIP datasets for some of these metrics, our findings point to specific settings and statistical parameters to improve differential HiChIP analysis. DiffHiChIP is publicly available at: https://github.com/ay-lab/DiffHiChIP.

## RESULTS

### DiffHiChIP: a comprehensive framework for detecting differential loops from HiChIP data

DiffHiChIP calls differential chromatin loops/interactions/contacts from HiChIP data between two conditions (e.g., disease vs control or between two different cell types) having one or more replicates (**Figure 1A**). DiffHiChIP supports both DESeq2 and edgeR as the underlying models for differential analysis. Specifically, for edgeR, DiffHiChIP includes both exactTest and GLM-based settings applied with different statistical tests (LRT or QLFTest). DiffHiChIP supports both Benjamini-Hochberg (denoted by **BH** in this manuscript) adjustment and independent hypothesis weighting (denoted as **IHW** in this work) to perform the multiple hypothesis testing correction of p-values and FDR control. For IHW, DiffHiChIP uses either the mean normalized counts across conditions *(baseMean*, recommended in^27,37^) for DESeq2, or log counts per million (*logCPM*) for edgeR, as the independent covariates. DiffHiChIP also implements a custom distance stratification using equal occupancy binning (**Methods**), inspired by FitHiC^35,36^, and provides a comprehensive comparison between these distance decay modeling techniques (**Figure 1A**). As both DESeq2 and edgeR rely on background count distributions for statistical modeling, DiffHiChIP further supports two different settings of background contacts for these models (**Figure 1A**). The first setting, denoted as the *complete background* or **A** for *all*, uses the union of nonzero HiChIP contacts (contact count > 0, significant or not) from all input samples. The second setting, denoted as the *filtered background* or **F**, employs the HiChIP contacts significant in at least one input sample, according to a user-defined FDR threshold *t* (default 0.1) for FitHiChIP^23^ calls. DiffHiChIP provides a comparative assessment between these background settings. If the custom distance stratification with equal occupancy binning is employed, corresponding settings are denoted by **A+D** and **F+D** for the complete and filtered backgrounds, respectively.

**Figure 1:**
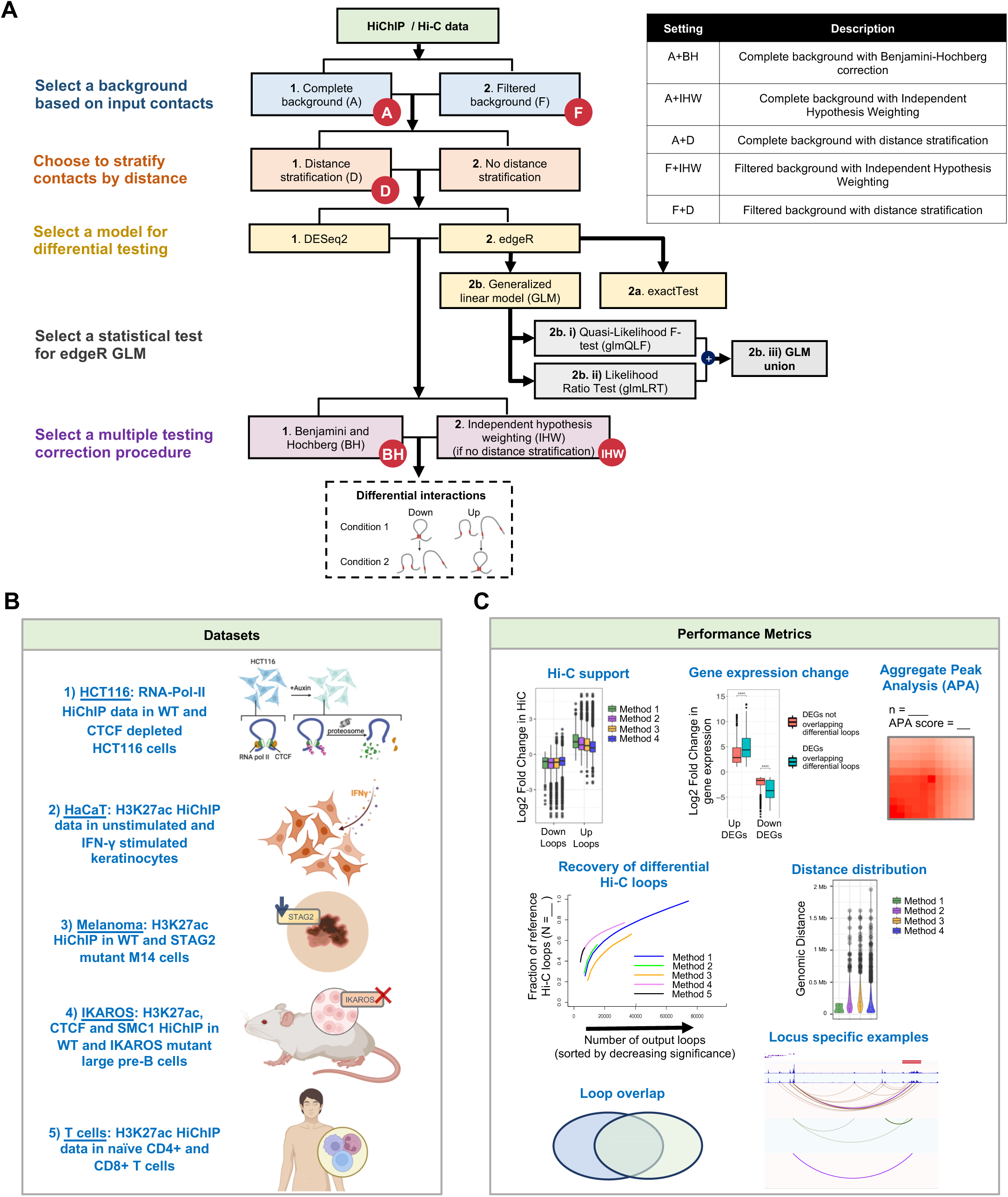
Workflow of DiffHiChIP: **(A)** DiffHiChIP calls differential loops from HiChIP data and supports various configurations related to: 1) employing complete (A) or filtered (F) background set of contacts, 2) choosing to utilize explicit distance stratification (D) or not, 3) using DESeq2 or edgeR, 4) for edgeR, using the exactTest or glm (LRT or QLFTest) techniques, 5) using either BH or IHW (except when distance stratification is used) for multiple testing correction. Table (right) summarizes all possible DiffHiChIP settings. **(B)** Schematic of the five different datasets employed in this study for performance validation (IKAROS has three different sets of HiChIP experiments). **(C)** Metrics employed to evaluate the differential loop calls reported by various settings of DiffHiChIP.

#### Description of datasets used for assessment of differential loop calls

We assessed DiffHiChIP and its various settings using five HiChIP datasets (**Figure 1B, Methods**): **1)** <u>HCT116:</u> RNA-Pol-II HiChIP from HCT116 colorectal cancer cells^14^ between control and Auxin treatment (CTCF depletion) conditions, **2)** <u>HaCaT:</u> H3K27ac HiChIP from keratinocytes (HaCaT) cells^15^ between wild-type (WT) and IFNγ stimulated (stim) conditions, **3)** <u>Melanoma:</u> H3K27ac HiChIP from M14 melanoma cells^13^ between wild-type (WT) and STAG2 knockdown (STAG2-KD) conditions, **4)** <u>IKAROS:</u> H3K27ac, CTCF and SMC1 HiChIP datasets from large pre-B cells in two conditions, namely wild-type (WT) IKAROS and DNA binding domain mutant IKAROS (IKDN)^40^, and **5)** <u>T</u> <u>cells</u>: H3K27ac HiChIP data of naïve CD4+ and CD8+ T cells from healthy blood donors. Datasets 1 to 4 have accompanying RNA-seq, Hi-C and ChIP-seq data for the corresponding conditions (except for dataset 3 – Melanoma, which lacks ChIP-seq), along with 2 HiChIP replicates per condition. The dataset 5, on the other hand, has 6 HiChIP replicates per condition. After preprocessing HiChIP datasets, we used FitHiChIP^23^ to call significant HiChIP loops (**Supplementary Data**, **Methods**) for different conditions and replicates, and used them as the inputs for DiffHiChIP. Differential loops are then evaluated using metrics derived from the input HiChIP data and other matching data available from the same conditions (**Figure 1C**).

### Comparison of different distance stratification approaches

Previous works such as HiCDC+^24^ binned chromatin loops per 10kb genomic distance and estimated DESeq2 size factors per bin, while another study^38^ computed the cumulative contact counts per 10kb distance bins and compared with the contact counts of the interactions having 140-150kb genomic distance. Both these techniques, however, did not capture long-range interactions. Thus, we implemented two new approaches to handle distance effect. The first approach adapts IHW by incorporating a covariate informative of the power of each test (ideally independent of p-values) in FDR control of p-values. We used the mean normalized counts (baseMean) for DESeq2 and log counts per million (logCPM) for edgeR as the independent covariates for IHW correction. The second approach implements a custom distance stratification (setting **D**) by adapting the *equal occupancy binning* technique implemented in our earlier work^23,36^. Here, chromatin contacts are stratified within a distance range (or bin) such that each bin would roughly have a similar number of contacts (**Methods**). Each range is then used separately as input to the chosen model for statistical significance estimation followed by BH correction across all bins for FDR control.

Next, using the complete background (setting **A**) as our starting point, we evaluated the overlap of DiffHiChIP loops reported by three different settings: 1) BH correction of p-values (**A+BH**) with no explicit distance correction, 2) IHW correction of p-values (**A+IHW**) with contact count-based covariates, and 3) equal-occupancy-based distance stratification (**A+D**) before significance estimation. Across all datasets, differential loops reported by A+BH were mostly the subsets of the corresponding loops from A+IHW (**Figure 2A, Supplementary Figure 1A-F**). Number of loops exclusive to the setting A+IHW were generally higher than those exclusive to A+D setting, except when using the edgeR glmLRT model, where the opposite was observed (**Figure 2A, Supplementary Figure 1A-F**).

**Figure 2.**
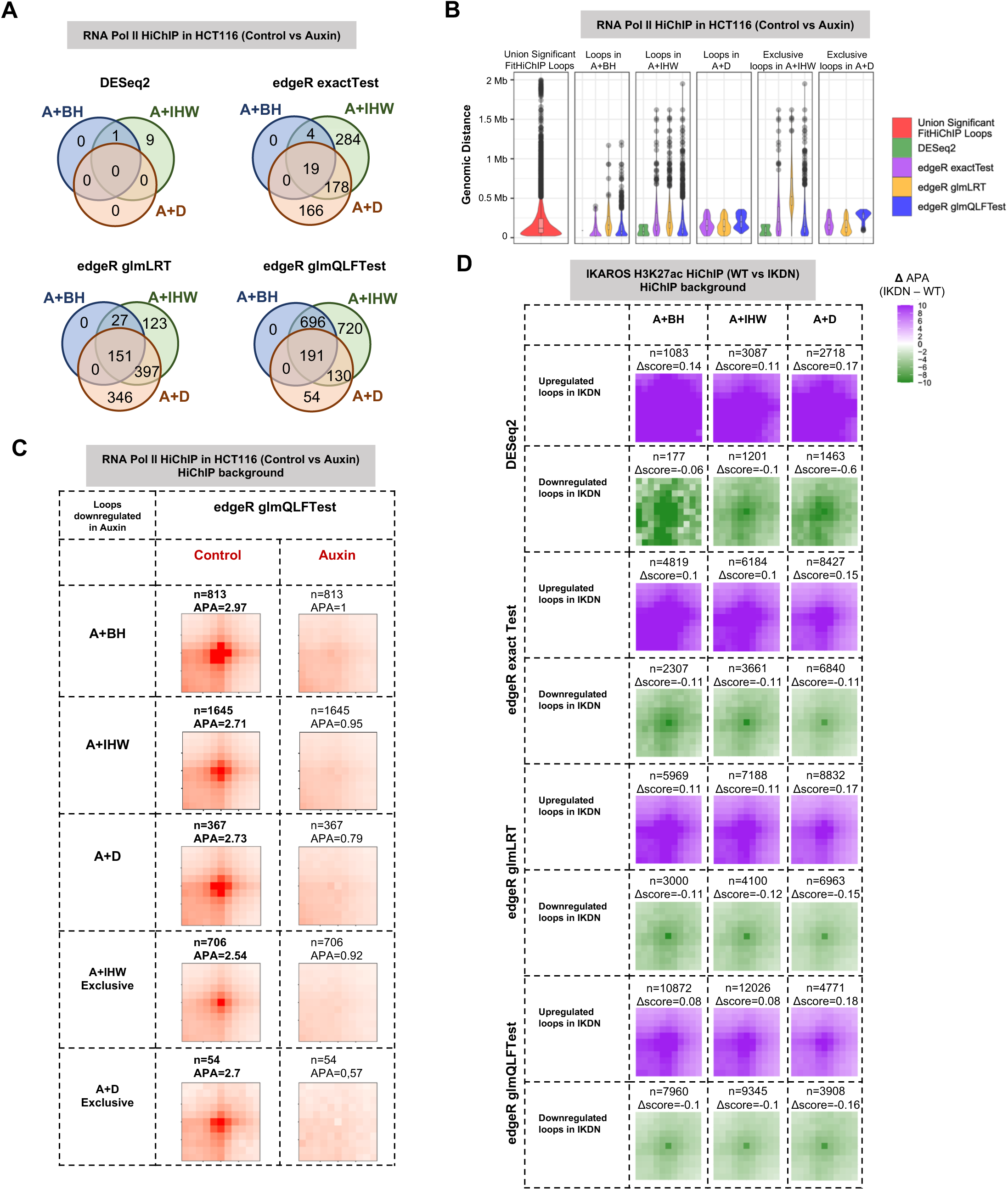
IHW correction detects longer-range differential loops better than BH or distance stratification (D): **(A)** Overlap of differential loops reported by A+BH, A+IHW and A+D settings for various DESeq2 and edgeR settings using the HCT116 cell line Pol II HiChIP data from wild type (control) and CTCF depleted (Auxin) conditions. DiffHiChIP is executed in the complete background (A) setting. **(B)** Genomic distance distributions for either all or exclusive A+IHW or A+D differential loops, along with the union of significant FitHiChIP loops of all replicates, for the same dataset as (A). **(C)** Aggregate peak analysis (APA) plots for the differential loop categories reported in (B) using HiChIP data as background. As majority of loop changes were downregulation/loss of loops upon CTCF depletion (Auxin), we only plotted results for this category with bold font indicating Control condition is where higher APAs are expected. The symbol “N” indicates the number of differential loops. **(D)** Differential APA plots (elementwise subtraction of the aggregate matrix for IKDN from that of WT) for IKAROS H3K27ac HiChIP data for various distance stratification settings. Differential APA scores (Δscore) represent the difference in APA scores between the IKDN and WT backgrounds.

#### Capturing long-range differential loops

We next assessed the performance of the A+IHW and A+D settings in modeling the distance decay of chromatin contacts. Differential loops from A+IHW particularly with edgeR settings included a subset of long-range loops, similar to overall set of significant loop calls, which were missing from A+D differential loops. For example, differential loops reported by A+D setting were shorter-range (distance < 400Kb) for HCT116 data (**Figure 2B**). Across different datasets, the upper quartile (75^th^ percentile) of the loop distance distribution for loops exclusively detected by A+IHW was higher (260Kb-1.2Mb) compared to those detected exclusively by A+D (90Kb-660Kb), suggesting that A+IHW captures longer-range differential loops more effectively (**Figure 2B, Supplementary Figure 1G-L**).

#### Support for differential loops by aggregate peak analysis of HiChIP data

We next employed aggregate peak analysis (APA)^2,23^ and differential APA (APA matrix of one condition subtracted from the other) to assess the relative enrichment of differential loops upregulated in specific conditions with respect to underlying HiChIP contact maps of the compared conditions (**Methods, Supplementary Figure 2A**), where higher magnitude of APA scores (or differential APA scores) indicate higher relative enrichment. Both the A+IHW and A+D settings, particularly when used with different edgeR configurations, reported similar APA (or differential APA) scores compared to the A+BH setting in spite of reporting much higher number of loops. In fact, loops exclusive to the A+IHW and A+D settings also showed higher APA (or differential APA) scores across different datasets (**Figure 2C-D, Supplementary Figure 2B-H**). In particular, loops exclusive to A+IHW setting showed some of the largest differential APA scores for IKAROS (pre-B cell) datasets between WT and DNA binding mutant IKAROS (IKDN) conditions (**Supplementary Figure 2F-H**). These results confirm the utility of IHW correction and distance stratification, compared to the classical BH adjustment of p-values.

#### Support for differential loops from analysis of matched Hi-C data

Availability of matched Hi-C data for the benchmarking studies prompted us to assess whether DiffHiChIP loops are also supported by differences in Hi-C signal between compared conditions. First, we performed differential APA analysis. For the loops downregulated upon CTCF depletion in HCT116 cells, we see strong differential APA patterns for A+BH, A+IHW and A+D for all different edgeR settings (**Figure 3A**). DESeq2 failed to report a sufficient number of loops to interpret any downstream analysis for the HCT116 data (**Figure 2A,3A**). Similar analysis for differential loops from all three IKAROS HiChIP experiments showed differential APA enrichment across all edgeR settings combined with either A+BH, A+IHW and A+D with H3K27ac HiChIP data showing the strongest enrichment scores (**Supplementary Figure 3A-C**). Next, for individual sets of differential loops detected from HiChIP data, we computed the log2 fold change of respective Hi-C contact counts between the compared conditions. Differences in Hi-C signal supported loss/decrease of looping for HiChIP differential loops detected by DiffHiChIP for the IKAROS and HCT116 datasets **(Figure 3B, Supplementary Figure 3D, 3G-I)**. We note that DESeq2 reported loops showed higher differences than those from edgeR settings only for the IKAROS H3K27ac HiChIP data (**Figure 3B, Supplementary Figure 3I**). Interestingly, the Hi-C fold change distributions were centered either near (Melanoma; **Supplementary Figure 3F**) or at zero (HaCaT; **Supplementary Figure 3E**) for two datasets suggesting that the differences in HiChIP signal were mainly related to changes in the underlying 1D signal or “loop visibility” rather than true changes in 3D organization. However, APA using HiChIP background supported the differential loop calls for these two datasets (**Supplementary Figure 2C-H, 6**) highlighting the difficulty of distinguishing true loop changes solely from differential HiChIP analysis. Lastly, to evaluate the support of DiffHiChIP loops in Hi-C data, we defined a stringent set of reference differential Hi-C loops by applying FitHiC2^36^ on the respective Hi-C datasets followed by a simple fold change criterion for filtering (**Methods**). We did not employ a specific statistical approach (such as DESeq2 or edgeR) to avoid any potential bias. The A+IHW and A+D settings particularly with edgeR glmQLFTest and glmLRT models, respectively, recovered higher fraction of reference Hi-C loops for IKAROS and HCT116 datasets (**Figure 3C, Supplementary Figure 4A, 4D-F**) whereas for HaCaT and melanoma datasets, edgeR glmLRT and exactTest performed similar and better than the glmQLFTest model **(Supplementary Figure 4B-C)**. Overall, adjustment of p-values using either IHW (A+IHW) or custom distance stratification (A+D) produced stronger Hi-C support. The underlying choice of statistical test mattered with A+IHW with glmQLFTest and A+D with glmLRT reaching highest levels of recovery in most cases. As previously discussed, A+IHW performed better than A+D in recovering longer-range loops and Hi-C data supported these A+IHW exclusive loops.

**Figure 3.**
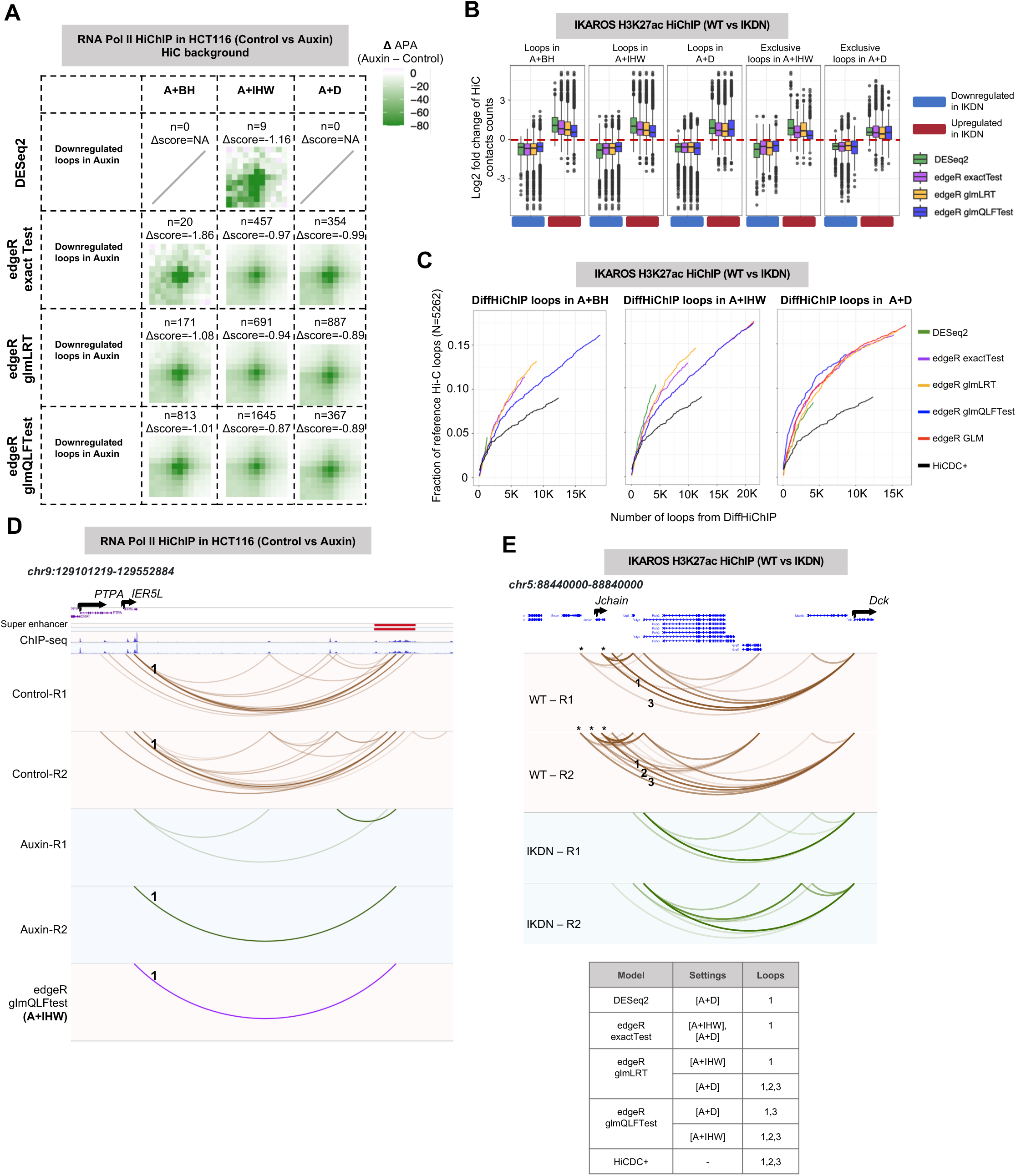
Further assessment of different distance stratification approaches: **(A)** Differential APA plots for HCT116 cell line Pol II HiChIP data for various distance stratification settings, using Hi-C data as background. Differential APA scores (Δscore) represent the difference in APA scores between the Auxin and Control backgrounds. **(B)** Log2 fold change (WT divided by STAG2-KD) in normalized Hi-C contact counts for upregulated and downregulated differential HiChIP loops for different settings of DiffHiChIP. We used IKAROS H3K27ac HiChIP data with or without IKAROS mutation. **(C)** Recovery of reference differential Hi-C loops by DiffHiChIP for different distance stratification settings and statistical tests for the dataset in (A). The plots also include HiCDC+ for comparison. X-axis shows the top-k number of loops called for each method and y- axis shows the fraction of recovered differential Hi-C loops. The symbol “N” indicates the number of reference Hi-C loops. **(D)** Differential loops lost upon CTCF depletion (Auxin) that linked the gene *IER5L* and the ∼400Kb downstream super enhancer (marked 1). This loop was detected as differential by only edgeR glmQLFTest for the A+IHW setting. **(E)** Three differential loops (marked 1, 2, 3; distance ∼250Kb) between WT and IKDN conditions for the IKAROS H3K27ac HiChIP data between *Jchain* and *Dck* genes, and their detection by different settings of DiffHiChIP and HiCDC+ represented in a tabular format.

#### Analysis of example loci for detecting long-range differential loops

To further compare different approaches in terms of their recovery of differences in long-range signals, we considered specific example loci analyzed in detail previously^14,40^. For HCT116 data, we focused on differential loops between *IER5L* and a ∼400kb downstream superenhancer (**Figure 3D**). Neither DESeq2 nor edgeR in A+H or A+D settings detected the ∼400kb loop downregulated upon CTCF depletion whereas A+IHW with glmQLFTest reported this loop as differential (**Figure 3D**). For the same dataset, we also looked at another loop connecting *MYC* and a ∼1.9Mb downstream superenhancer near the gene *GSDMC*, which was also detected only by A+IHW with glmQLFTest (**Supplementary Figure 4G**). For IKAROS H3K27ac HiChIP data, we focused on the *Jchain* locus, where we reported loss of >250kb loops in our previous work ^40^. Consistent with higher recovery in genome-wide results **(Figure 3C)**, A+IHW with glmQLFTest and A+D with glmLRT were the two combinations that captured differences in all three of the indicated loops whereas other settings missed either one or two of them **(Figure 3E).** These results support the above-mentioned genome-wide observations in terms of A+IHW setting’s superiority for capturing *bona fide* long-range differences.

### edgeR with glm models provides higher recall for differential loop calling

We next assessed DESeq2 and various statistical tests from edgeR (exactTest, glmLRT and glmQLFTest) for differential loop calling when coupled with the A+IHW setting. DESeq2 reported a lower number of differential loops, most of which were covered by edgeR glm settings for all datasets (**Figure 4A, Supplementary Figure 5A-F**). Among the two glm models, glmQLFTest reported a much higher number of differential loops in most cases (4 out of 6) compared to glmLRT. For the other two cases (HaCaT and Melanoma), glmLRT and edgeR exactTest reported highly overlapping differential calls that are missed by the glmQLFTest (**Supplementary Figure 5B-C**). This prompted us to define another set of differential loops named **edgeR GLM** to denote the union of loops from glmLRT and glmQLFTest. Although APA between these glm models alongside DESeq2 and edgeR exactTest did not reveal any striking difference in their APA scores, loops from DESeq2 showed stronger patterns in the bottom left portion (i.e., the area that remains between the two anchors) highlighting the dominance of shorter-range loops among those reported by DESeq2 (**Figure 4B, Supplementary Figure 6**). Considering the recovery of reference Hi-C loops, edgeR glmQLFTest settings recovered higher overall fraction of loops for 4 out of 6 datasets (same 4 as above), but glmLRT settings led to a better ranking of significance evidenced by higher recovery at an equal number of differences reported (i.e., same value on the x-axis) (**Supplementary Figure 4A-F**). The edgeR exactTest and glmLRT models performed similarly for the remaining two datasets where glmQLFTest reported a low number (**Supplementary Figure 4B)** or no differential loops (**Supplementary Figure 4C)**. HiCDC+ reported very low recovery across all conditions (**Supplementary Figure 4A-F**).

**Figure 4.**
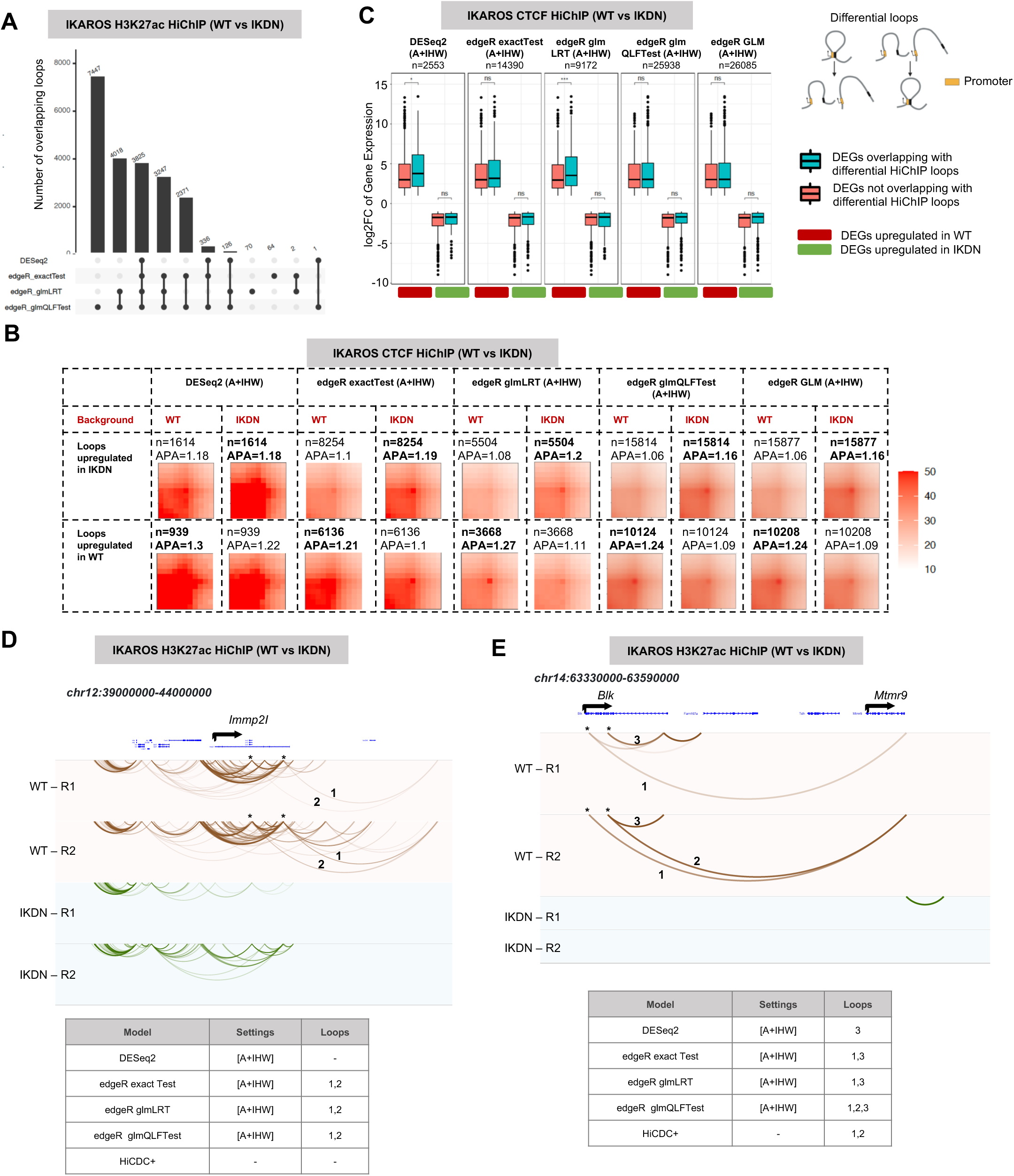
edgeR with GLM setting provides higher recall for differential loop calling: **(A)** Overlap of differential loops between DESeq2 and various edgeR settings for the complete background with IHW corrected FDR (setting A+IHW) for IKAROS H3K27ac HiChIP data. **(B)** APA plots for IKAROS CTCF HiChIP data between WT and IKDN conditions, for differential loops from DESeq2 and different edgeR settings. Values in bold denote expectation of higher APAs among the two conditions (i.e., upregulated loops in that conditions. **(C)** Enrichment of magnitude of gene expression change (log2 fold change) for differential genes segregated with respect to their overlap with DiffHiChIP loops from different settings for the IKAROS CTCF HiChIP dataset. Enrichment is computed separately for genes upregulated in either condition (WT or IKDN). The symbol “n” indicates the number of differential loops in different categories. Model (top right) of differential loops that overlap a gene promoter in at least one anchor. Significance was calculated using a Wilcoxon test (two-sided). *P<=0.05; **P<=0.01; ***P<=0.001; ****P<=0.0001; ns, not significant. **(D)** Example of two differential loops (∼2Mb) in the *Immp2I* locus lost upon loss of IKAROS (IKDN) for the H3K27ac HiChIP dataset. The associated table shows recovery of corresponding loops with different settings of DiffHiChIP and with HiCDC+. **(E)** Three differential loops for *Blk* locus lost upon loss of IKAROS and their tabulation, similar to panel D. Loops 1 and 2 have ∼170Kb distance.

Our previous study showed that differences in regulatory HiChIP contacts are associated with larger changes in the expression of genes at the loops anchors^11^. Thus, we assessed the association between fold changes of differentially expressed genes (DEGs) and presence of differential HiChIP loops from DiffHiChIP across different settings (**Methods**). Our results indicate that, in many cases, there is a statistically significant difference between the fold changes of DEGs overlapping with differential HiChIP loops compared to those that do not (**Figure 4C, Supplementary Figure 7**). Looking at differences across methods, this difference was generally more prominent for DESeq2, edgeR exactTest and glmLRT differential loops although the results highly varied across datasets (**Figure 4C, Supplementary Figure 7**).

Lastly, we considered specific genomic loci previously reported to have long-range differential loops. One example is the *Immp2I* locus (∼5Mb shown) and two long-range (∼2Mb) H3K27ac HiChIP loops that were present in WT but lost in the IKDN condition compared^40^. We observed that all edgeR settings retrieved these two loops as differential but DESeq2 and HiCDC+ did not (**Figure 4D**). We also considered the ∼250Kb *Blk* locus and WT specific H3K27ac loops that link *Blk* to distal enhancers and promoters. Out of three highlighted loops, DESeq2 reported only the short-range one as differential while edgeR exactTest and glmLRT reported both the short range and one long range loop that was consistently significant across the two WT replicates (**Figure 4E**). On the other hand, HiCDC+ captured both long range loops but missed the short-range loop that was reproducibly observed across replicates. glmQLFTest reported all three of the highlighted loops as differential in the A+IHW setting. Combined with the three other examples discussed above (**Figure 3D-E, Supplementary Figure 4G**), these results highlight the importance of using edgeR glm-based model for increased sensitivity.

### Comparison of different background estimation options

All the results so far used the complete background (A), basically union of HiChIP contacts (significant or not) across all input samples as the background for underlying DESeq2 or edgeR settings. Previous studies such as HiCDC+^24^ and our previous work FitHiChIP^23^, on the other hand, used a filtered subset of chromatin contacts to infer a background by considering only the contacts having FDR < *t* (user-defined threshold) in at least one input sample. FitHiChIP^23^ used *t* = 0.01 while HiCDC+ employed a more lenient *t* = 0.1. This approach of performing background estimation from only strong contacts (or loops) eliminates insignificant contacts with either very low contact counts, or contacts having short genomic distance and high (but not statistically significant) contact counts, likely reducing some of the false positive discoveries from a more lenient background. We implemented a similar filtered background estimation for DiffHiChIP which considers union of loops calls from the compared conditions (default *t* = 0.1) and coupling of this with two different distance stratification methods: 1) filtered background with IHW (model **F+IHW**), 2) filtered background with distance stratification (model **F+D**).

First, we compared the complete (A+IHW) and filtered (F+IHW) backgrounds for IHW. For the HCT116 and IKAROS HiChIP datasets, loops from the setting A+IHW mostly included the loops from the setting F+IHW with ∼1.5 to 3 times more loops except the HaCaT and Melanoma datasets (**Figure 5A, Supplementary Figures 8A-F**). However, the F+IHW setting did not lead to any noticeable improvement in APA enrichment scores neither for IKAROS nor for any other dataset (**Figure 5B, Supplementary Figures 10,11**), even though for CTCF HiChIP data the F+IHW setting had ∼60% reduction in differential loops compared to A+IHW (**Figure 5B).** Considering the recovery of differential Hi-C loops, we mainly observed a decrease in overall recovery by F+IHW with some exceptions such as when edgeR glmLRT model is used (**Figure 5C, Supplementary Figures 8G-L**). We also assessed the fold change of differentially expressed genes when they overlap with differential loops from F+IHW setting (**Figure 5D, Supplementary Figure 9**) and compared it to A+IHW, but did not observe any noticeable or generalizable pattern of higher enrichment between A+IHW and F+IHW (**Supplementary Figure 7** vs **9**).

**Figure 5.**
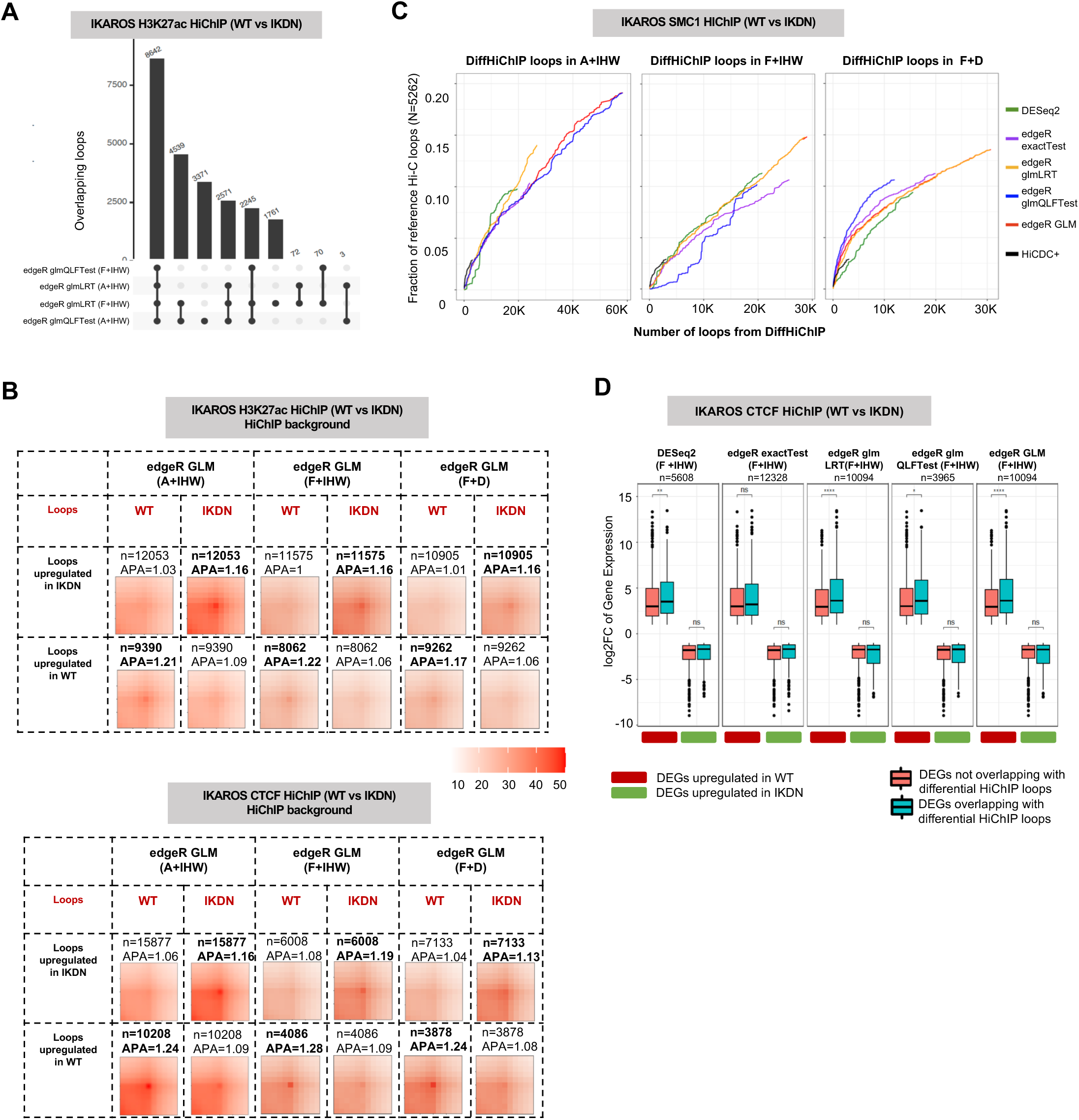
Filtered background produces highly enriched differential loops: **(A)** Overlap of differential loops for various edgeR glm settings between the complete (A+IHW) and filtered (F+IHW) background settings, for the IKAROS H3K27ac HiChIP data. **(B)** APA plots for edgeR GLM setting (union of LRT and QLFTest) and for either A+IHW or F+IHW settings, with respect to IKAROS H3K27ac (top) and IKAROS CTCF (bottom) HiChIP datasets. Values in bold denote expectation of higher APAs among the two conditions (i.e., upregulated loops in that conditions). **(C)** Comparison between the complete (A) and filtered (F) backgrounds (A+IHW, F+IHW and F+D) with respect to their recovery of differential Hi-C loops by different settings of DiffHiChIP for IKAROS SMC1 HiChIP dataset. The symbol “N” indicates the number of reference Hi-C loops. **(D)** Enrichment of magnitude of gene expression change (log2 fold change) for differential genes segregated with respect to their overlap with differential loops from different settings of DiffHiChIP executed with the F+IHW setting for the IKAROS CTCF HiChIP dataset. Enrichment is computed separately for genes upregulated in either condition (WT or IKDN). The symbol “n” indicates the number of differential loops in different categories. Significance was calculated using a Wilcoxon test (two-sided). *P<=0.05; **P<=0.01; ***P<=0.001; ****P<=0.0001; ns, not significant.

Comparing between IHW and distance stratification (**D**) using the filtered background (F+IHW and F+D) across all edgeR and DESeq2 settings and all datasets, F+IHW led to slightly higher APA scores for most cases but this was not always the case and the differences were minimal (**Figure 5B, Supplementary Figures 10,11**). The F+IHW setting, particularly using edgeR models, recovered higher fraction of reference Hi-C loops than the F+D setting in some datasets and showed comparable performance for the remaining (**Figure 5C, Supplementary Figures 8G-L**).

Overall, these results suggested limited utility of filtered background in terms of decreasing potential false positive calls. Although they reported a similar number of differential loops, filtered background coupled with IHW correction generally performed better than distance stratification (F+D), especially in recovering differential Hi-C loops.

### DESeq2 sensitivity increases with higher number of replicates

Most of the HiChIP datasets in reference studies (and the results used so far) have very few (either 1 or 2) replicates per condition. To assess various settings of DiffHiChIP, we next turned to a HiChIP dataset with a higher number of replicates. We used our previously published HiChIP data^11^ for naïve CD4 and naïve CD8 cell types each with six different donor samples as “replicates”. Such higher number of replicates considerably increased the number of differential loops reported by DESeq2 (A+IHW) with ∼5 times more differential loops compared to various edgeR settings (**Figure 6A**) with a similar genomic distance distribution (**Figure 6B**). These results suggest that the output of DESeq2 is highly dependent on the number of replicates, an observation previously reported with respect to differential analysis of RNA-seq data^41^. When the DESeq2 and edgeR results are compared with respect to different background choices, edgeR models reported a higher number of loops and lower APA enrichment in the filtered (F+IHW) setting compared to A+IHW (**Figure 6C**). Given the common T cell lineage and naïve state of CD4+ and CD8+ T cells, the aggregate analysis of the reported differential loops did not lead to strong APA enrichment in any setting (edgeR or DESeq2). Lack of any visible differential enrichment in the upregulated loops for most of the DESeq2 results together with high sensitivity to background choice suggest a large number of false positives (**Figure 6C**). When we specifically looked at differential loops involving the loci containing *CD8A* and *CD8B* (markers of CD8+ T cells), these were detected similarly by most edgeR settings and DESeq2 (**Figure 6D**) as upregulated in CD8+ T cells as expected. Further comparative analysis of different datasets with large number of replicates (>3) from a case/control experiment setting (e.g., CTCF or cohesin depletion) are needed to fully characterize the extent of increased sensitivity of DESeq2 and the portion of true positive versus false positive discoveries that come with this increase.

**Figure 6.**
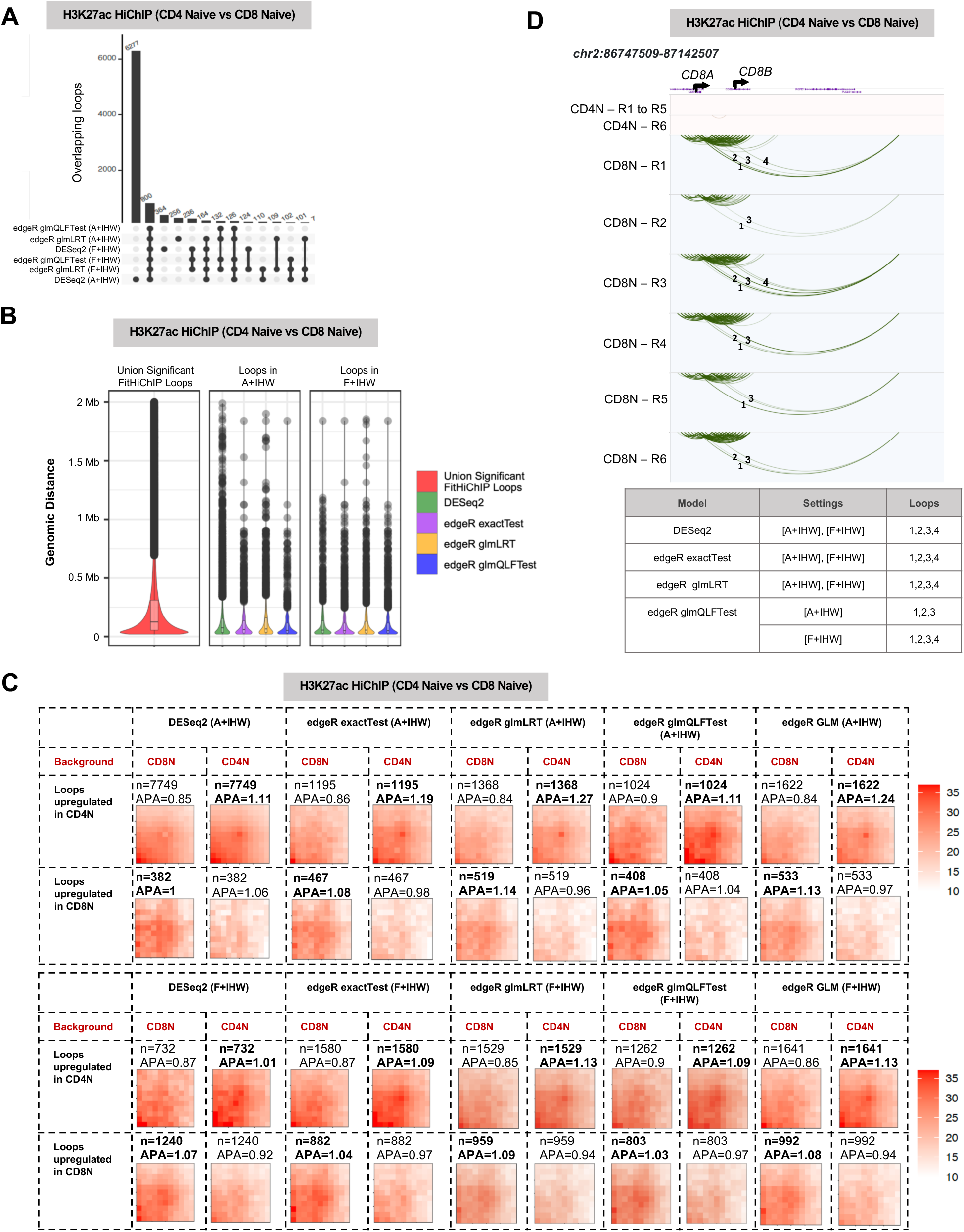
DESeq2 is highly sensitive for higher replicates and background models: **(A)** Overlap of differential loops between naïve CD4 and naïve CD8 cell types for various settings of DiffHiChIP with respect to either settings A+IHW or F+IHW. **(B)** Genomic distance distributions for all A+IHW or F+IHW differential loops, together with the union of significant FitHiChIP loops of all replicates. **(C)** APA plots for DESeq2 and different edgeR settings and for the models A+IHW and F+IHW. Values in bold denote expectation of higher APAs among the two conditions (i.e., upregulated loops in that conditions). **D)** Differential loops between naïve CD4 and naïve CD8 in the *CD8A* locus for different settings of DiffHiChIP, with respect to the backgrounds A+IHW and F+IHW. Four long-range loops involving the genes *CD8A* and *CD8B* are indicated by numbers.

## DISCUSSION

Recent progress of NGS-based 3C technologies, particularly HiChIP, enabled researchers to understand the gene regulation by *cis* regulatory elements (e.g., enhancers) via 3D chromatin loops^42^. Differences in DNA looping structure are associated with variation in gene expression and cell states. Thus, chromatin loops differential between two conditions (disease versus healthy, treatment conditions, or cell types) reveal the underlying regulatory changes and corresponding genomic segments. Although there are methods developed by us and others for identifying differential HiChIP loops^23–25^, as well as for Hi-C and PCHi-C data^19–22^, metrics to compare their results and studies that do this systematically across distinct datasets are lacking. Also, since the difference in HiChIP looping may be due to the changes in either chromatin folding or the underlying 1D (ChIP-seq) distribution, this assessment is even harder for HiChIP data. To date, no study has benchmarked differential HiChIP loop callers and assessed the impact of different distance stratification methods, background estimation and statistical tests employed on their performance. Here we present the first such large-scale benchmarking study while introducing new approaches (e.g., distance stratification) and creating a codebase that implements and makes available all the evaluated approaches for differential HiChIP loop calling.

DiffHiChIP is a comprehensive framework which incorporates reference count-based models DESeq2 and edgeR exactTest using either complete or pre-filtered backgrounds, includes edgeR-based glm models, and simultaneously supports distance stratification by IHW and custom implementation. GLM-based regression coupled with LRT or QLFTest are expected to model the higher dispersion of HiChIP contacts better, while IHW-adjusted p-values model the distance decay of chromatin contacts. DiffHiChIP is the first approach incorporating edgeR glm models for differential HiChIP analysis motivated by their application in differential Hi-C loop calling (diffHiC^43^) and in single-cell RNA-seq studies modeling differential abundance^33,34^.

Although DESeq2 also computes regression by glm, it estimates gene-wise dispersions and uses the Wald test for significance estimation. The glm models in edgeR support both common and gene-wise dispersions, and the LRT or QLFTest are more reliable for modeling non-linear decay of HiChIP contacts with higher dispersion compared to 1D RNA-seq or ChIP-seq datasets. We note that these glm models in DESeq2 or edgeR are, however, applicable when both input conditions have at least two replicates. When only a single replicate is available, DiffHiChIP defaults to applying the edgeR exactTest setting.

Our results show that classical BH-adjusted p-values miss out on the differences of long-range chromatin interactions due to their lower contact counts, while the IHW or custom distance stratification techniques better recover them by modeling their distance decay. The IHW correction particularly performs well in capturing differences in long-range loops evidenced by multiple lines (different metrics) of support for such differences. For IHW, we did not use genomic distance as the covariate (as suggested in^38^) since the *baseMean* or *logCPM* values adequately represent the distance decay of chromatin contacts.

Higher number of replicates (>3 per condition) increases the number of detections by DESeq2, a phenomenon highlighted in a previous study^41^ which also suggested using simple Wilcoxon rank-sum tests for count-based RNA-seq datasets with high number of replicates. However, from our analysis, it was not clear what fraction of the additional discoveries by DESeq2 were *bona fide* changes and not false positive. Given that HiChIP datasets usually have lower number of replicates (1 or 2) per category, edgeR models with IHW are potentially more preferable given their applicability to most cases and superior performance according to multiple different metrics.

DiffHiChIP also incorporates options from existing differential HiChIP loop callers diffloop and FitHiChIP which employ edgeR exactTest with pre-filtered background, with diffloop additionally filtering out loops detected in a single sample. Another published method HiCDC+ is employs DESeq2 with library size factors estimated separately for individual distance bins (default 10Kb). Our results show that HiCDC+ and DESeq2 recover lower fraction of reference Hi-C loops and do not detect differential loops in various example loci, compared to the edgeR glm models suggesting lower sensitivity. Similar to our earlier work FitHiChIP^23^, DiffHiChIP can identify the differential and non-differential loop anchors between conditions by applying edgeR onto the 1D ChIP-seq (if additionally provided as an input) coverage between conditions, and characterize the differential interactions involving non-differential HiChIP loop anchors and thus are explained largely by the differences of 3D chromatin folding between conditions.

It is important to note that none of the specific metrics we employed are informative of superior performance on their own and have to be represented in the context of all other metrics. For instance, APA scores could be maximized by capturing a very small number of loops with largest differences but this will lead to low sensitivity. It is also possible that higher APA for one method compared to the other can come at the cost of the first method missing out on longer range loops. Better recovery of differential Hi-C loops by one particular method may be mostly a result of a much larger number of differential calls that may be related to lower specificity. Therefore, we chose to present all these different metrics alongside one another, which, admittedly complicated our presentation.

Overall, DiffHiChIP is a comprehensive framework for differential HiChIP analysis that combines multiple approaches used to date and introduces new options to improve capture of differences in long-range loops alongside shorter range loops. With the ever-increasing number of HiChIP datasets generated to compare multiple conditions/perturbations across different biological systems, we believe the presented results will be of high interest to the field and the developed framework will be highly utilized. We make our documented source code and package available on GitHub and all of the produced data files (differential loop calls across all datasets and all DifHiChIP setting discussed in this work) through a webserver at https://ay-lab-tools.lji.org/DiffHiChIP/

## MATERIALS AND METHODS

### Dataset description

We used the following HiChIP datasets to validate DiffHiChIP**: 1)** RNA-Pol-II HiChIP data from HCT116 colorectal cancer cells^14^ (available from Gene Expression Omnibus or GEO repository under the accession number GSE179545) in two conditions: untreated (or control) and treated with Auxin to induce a degron system on CTCF. This dataset has Hi-C, RNA-seq and RNA-Pol-II ChIP-seq data for the corresponding conditions and replicates. **2)** H3K27ac HiChIP data in unstimulated and IFN-γ stimulated HaCaT cells^15^ (GEO: GSE151193). Accompanying RNA-seq and Hi-C data for the corresponding conditions are also provided. **3)** H3K27ac HiChIP data from M14 melanoma cells expressing doxycycline-inducible shRNA targeting STAG2 treated with (STAG2-KD) or without (WT) doxycycline^13^ (GEO: GSE156773). **4)** H3K27ac, CTCF and SMC1 HiChIP from mouse large pre-B cells in two conditions: wild-type (WT) and IKAROS mutant (IKDN), which correspond to an *in-vivo* deletion of *Ikzf1* exon 5 encoding the IKAROS DNA-binding domain^40^ (GEO: GSE232490). This dataset has matching Hi-C, RNA-seq and H3K27ac, CTCF and SMC1 ChIP-seq data. **5)** H3K27ac HiChIP data in two immune cell types naïve CD4 and naïve CD8 prevalent in human peripheral blood mononuclear cells (PBMCs) from six donors^11^. This data is available through the database of Genotypes and Phenotypes (dbGaP) under the accession number phs001703.v3.p1. We note that datasets 1 to 4 have 2 replicates per condition, while the dataset 5 has 6 replicates (different donors) per condition.

### HiChIP data processing and loop calling

HiChIP paired-end reads were aligned to the human hg38 or mouse mm10 (for the IKAROS dataset) genome assembly using the HiC-Pro^44^ pipeline (version 2.11.4). Default settings were used to remove duplicate reads, assign reads to restriction fragments, filter for valid pairs, and generate raw and ICE normalized interaction matrices at a range of resolutions. For visualization, valid pairs were converted to .hic files using the script *hicpro2juicebox.sh* from Juicebox v3.1.0^45^. ChIP-seq peak calling was performed using MACS2^46^ (version 2.2.9.1) with input chromatin as control and with a q-value cutoff of 0.05. FitHiChIP^23^ (version 9.1) was used to identify statistically significant HiChIP interactions from individual samples, employing the ChIP-seq data generated independently from the same study/conditions as the HiChIP libraries. HiChIP interactions were called using peak-to-all background setting (UseP2PBackgrnd = 0), 5Kb bin size, and distance range between 10Kb - 3Mb. For the SMC1 and CTCF HiChIP datasets in^40^, only peak-to-peak interactions were assessed for significance, i.e., when both anchors overlapped a ChIP-seq peak. For all other HiChIP datasets, peak-to-all interactions were considered, i.e., at least one anchor needed to overlap with a reference ChIP-seq peak.

### Overview of DiffHiChIP

#### Input data

DiffHiChIP is a comprehensive framework for calling differential loops primarily from HiChIP data. It employs loop calls derived for the input samples computed using either FitHiChIP^23^ or other HiChIP loop callers. We note that all input samples should be processed by the same chromatin loop caller employing the same parameters (resolution, distance thresholds, etc.). Complete list of all interactions for individual samples (whether they are loops with respect to some filtering by their significance values) along with their statistical significance values are provided as an input to DiffHiChIP.

#### Background loops

DiffHiChIP supports two different sets of background loops for the underlying DESeq2 or edgeR settings: 1) complete (A) background: using the union of chromatin interactions (nonzero contact counts) from all the input samples, and 2) filtered (F) background: using the union of loop calls with FDR < *t* for at least one input sample, where *t* is a user-defined FDR threshold with default 0.1 (similar to HiCDC+^24^).

#### Significance thresholds for differential analysis

After applying the DESeq2 or edgeR models for significance estimation, loops with adjusted p-values (from DESeq2 or edgeR models) < *f*, absolute log fold change > *l,* and statistically significant (FDR from the chromatin loop caller < *t*) in at least one input sample are returned as differential, where *f*, *l* and *t* are user-defined thresholds with default values of 0.05, 1 and 0.01, respectively.

#### Applying DESeq2

The design variable for DESeq2 is constructed using the input condition information. Reference functions from the DESeq2 Bioconductor package such as *DESeqDataSetFromMatrix*, *DESeq*, and *results* are employed.

#### Applying edgeR

DiffHiChIP incorporates edgeR supporting both exactTest and GLM models. Reference routines from the edgeR Bioconductor package are used, such as *estimateDisp* for estimating dispersions, *exactTest* for the exactTest model, *glmFit* and *glmLRT* for GLM with LRT model, and *glmQLFit* and *glmQLFTest* for modeling GLM with QLFTest.

#### IHW for distance stratification

DiffHiChIP supports applying independent hypothesis weighting (IHW) on the resulting p-values from DESeq2 or edgeR, by applying the routine *ihw* from the Bioconductor package IHW. The baseMean and logCPM values for each interaction are used as the covariates for DESEq2 and edgeR, respectively. The *alpha* parameter for *ihw* routine is kept the same as the user-defined significance threshold *f* for differential analysis (mentioned above).

#### Custom distance stratification

DiffHiChIP also provides a custom implementation of distance stratification of chromatin loops, to mitigate the distance decay bias. We adapted the equal occupancy binning described in our previous method FitHiChIP^23^. If *N* is the number of locus pairs and *C* is the total number of contacts between them (sum of contact counts), then considering *M* bins (we considered *M* = 300 for the default distance range from 10 Kb to 3 Mb equally spaced by 10 Kb), each bin would have ∼ *C* / *M* contacts. We first sorted the interactions by their genomic distance values, assigned them into 10 Kb binning intervals according to their interaction distance values, and then constructed the equal occupancy bins such that each bin has at least *C* / *M* contacts. Each of these equal occupancy bins and their constituent interactions were then applied to the downstream DESeq2 or edgeR models for estimating the p-values. Finally, all the p-values from all the interactions across all equal occupancy bins are subjected to BH-correction. Note that when this distance stratification is employed, IHW is not used to avoid double correction.

### Aggregate peak analysis

To show the average contact count distribution of loops and their surroundings, we performed an aggregate peak analysis (APA) of loop calls using the R package GENOVA v1.0.0.9^47^. For aggregate signal assessment, we used either HiChIP or Hi-C contact maps binned at 10 kb resolution that were normalized by Knight-Ruiz (KR). We retrieved the normalized contact counts of a 100kb x 100kb region centered on each loop coordinate corresponding to a pair of loci on the same chromosome represented by coordinates: *i* and *j*. Without loss of generality, we can assume coordinate *i* < *j* and represent their genomic distance by d=d(*i*,*j*)=*i* – *j* for a given loop. The APA then plots the average contact count across all 100kb x 100kb with center pixel depicting (*i*,*j*), bottom left pixel corresponding to (*i*+50kb, *j*-50kb) and top right pixel denoting (*i*-50kb, *j*+50kb) with genomic distances of d, d-100kb and d+100kb, respectively. In alignment with the literature, only loops with a genomic distance greater than 130kb are considered in the APA analysis. The APA score displayed on top of each plot is the ratio between the central pixel value and the mean value of pixels 15-30 kb downstream of the upstream loci and 15-30 kb upstream of the downstream loci.

We computed APA scores separately for the loops upregulated in each condition and, for each set, we use the underlying contact map from each condition separately, thus, providing us with four different APA plots and scores. Let *S_XY_* be the APA score for the loops upregulated in the condition *X* and with respect to the background set of HiChIP contacts from the condition *Y*. A set of differential loops between two input conditions *A* and *B* should ideally satisfy the condition *S_AA_ >= S_AB_* and *S_BB_ >= S_BA_*, that is, differential loops upregulated in a given condition should also be more enriched in the respective background set of loops. We denote *d_AB_* = (*S_AA_* - *S_AB_*) as the *differential APA score* for the loops upregulated in the condition *A*. Thus, higher values of *S_AA_* and *d_AB_* indicate higher enrichment of loops in the condition *A*.

### Calling superenhancers from ChIP-seq peaks

For the Auxin dataset (GEO: GSE179545) we identified superenhancers with the Rank Ordering of Super-Enhancers (ROSE) v1.3.2 algorithm^48^ using the H3K27ac ChIP-seq peaks and the default stitching size of 12.5 kb.

### Overlap of HiChIP / Hi-C loops between different models

To compare HiChIP loops between different settings of DiffHiChIP, we used the exact overlap strategy (i.e., identical anchors). To check the recovery of reference differential Hi-C loops by the DiffHiChIP output loops, we employed a slack of 5Kb similar to our previous work^23^, that is, declared two different loops overlapping if their respective anchors were within 5Kb (one bin size) of each other.

### Recovery of differential Hi-C loops by DiffHiChIP

Hi-C paired-end reads were aligned to the human hg38 or mouse mm10 (for the IKAROS dataset) genome assembly using the HiC-Pro v2.11.4^44^ pipeline as described under the HiChIP data processing section. FitHiC2^36^ was used with default parameters to identify statistically significant Hi-C interactions. We defined the reference differential Hi-C loops using EdgeR_exactTest and a criterion of log fold change > 2 between the input conditions. We then computed the fraction of these reference differential Hi-C loops recovered by the differential HiChIP loops for different settings of DiffHiChIP, subject to increasing number of loop calls (decreasing stringency), and according to the above-mentioned criteria of loop overlap.

### Differentially expressed genes and their overlap with differential loops

RNA-seq paired-end sequencing reads were aligned to the human hg38 or mouse mm10 (for the IKAROS dataset) genome assembly using STAR v2.7.1^49^ with default parameters. Normalization and differential gene expression analysis was performed using DESeq2 v1.40.2^27^. Differentially expressed genes (DEG) were identified using an adjusted p-value cutoff of 0.05. A DEG was considered to overlap with differential HiChIP loops if its transcription start site (TSS) was located within a differential loop anchor. The overlap of genomic regions was performed using the R package GenomicRanges v1.42.0.

### Statistics and reproducibility

R v4.0.1 was used for statistical analysis and plotting of data. The statistical significance between two groups was assessed using a two-sided Wilcoxon test.

### Code availability

DiffHiChIP is publicly available in the GitHub repository https://github.com/ay-lab/DiffHiChIP

## Data availability

### Supplementary Data 1

Lists the input HiChIP datasets, HiChIP statistics, and the number of FitHiChIP loops for these datasets.

### Web server

Differential loops for various datasets and for different settings of DiffHiChIP, and corresponding WashU browser tracks are hosted in the web server https://ay-lab-tools.lji.org/DiffHiChIP/

