## Supplementary Figures for "DiffHiChIP: Identifying differential chromatin contacts from HiChIP data"

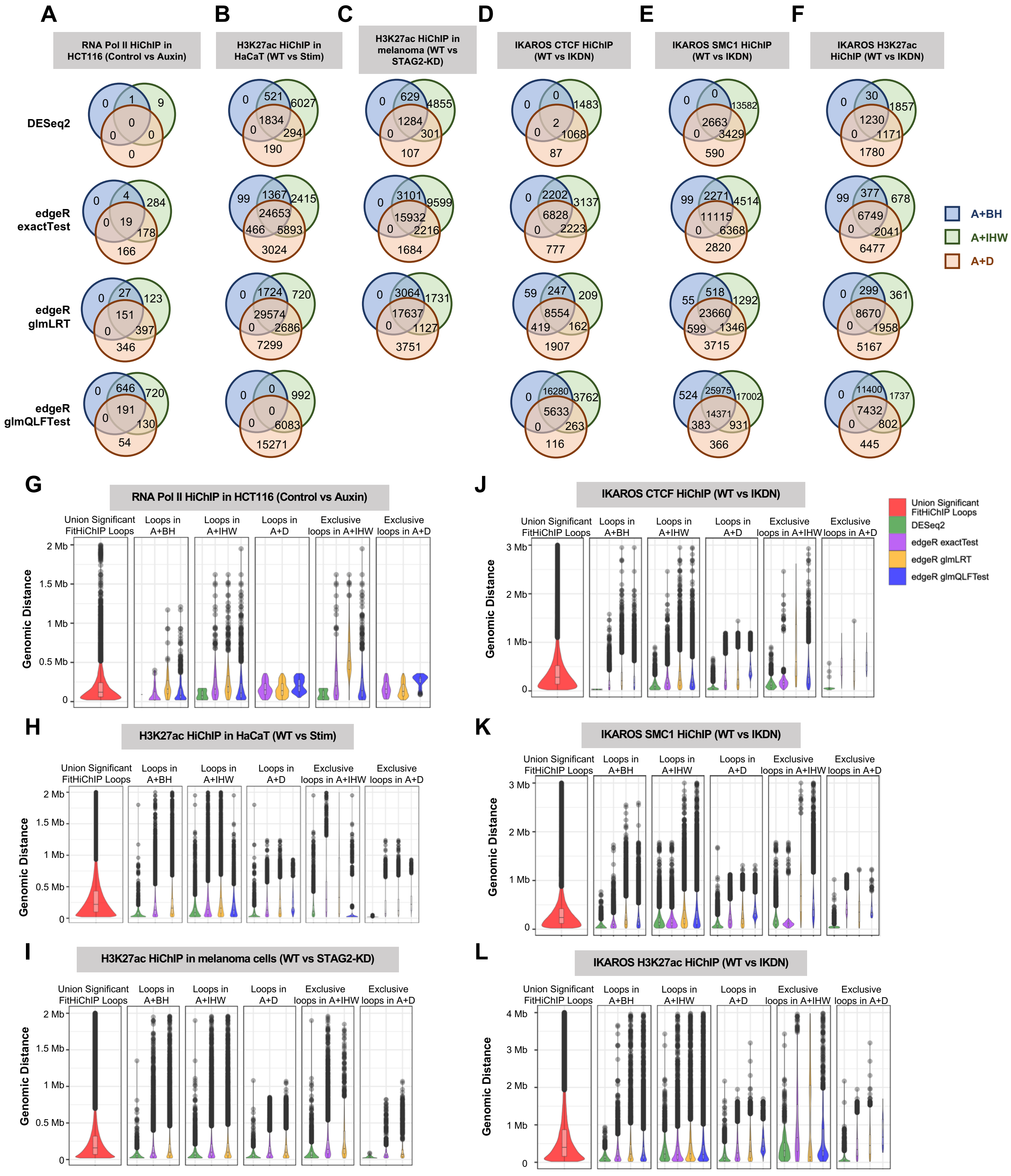

A
RNA Pol II HiChIP in HCT116 (Control vs Auxin)
B
H3K27ac HiChIP in HaCaT (WT vs Stim)
C
H3K27ac HiChIP in melanoma (WT vs STAG2-KD)
D
IKAROS CTCF HiChIP (WT vs IKDN)
E
IKAROS SMC1 HiChIP (WT vs IKDN)
F
IKAROS H3K27ac HiChIP (WT vs IKDN)
DESeq2
edgeR exactTest
edgeR glmLRT
edgeR glmQLFTest
0 1 9 0 0 0 0
0 521 6027 1834 0 294 190
0 629 4855 1284 0 301 107
0 0 1483 2 0 1068 87
0 0 13582 2663 0 3429 590
0 30 1857 1230 0 1171 1780
0 4 284 19 0 178 166
99 1367 2415 24653 466 5893 3024
0 3101 9599 15932 0 2216 1684
0 2202 3137 6828 0 2223 777
99 2271 4514 11115 0 6368 2820
99 377 678 6749 0 2041 6477
0 27 123 151 0 397 346
0 1724 720 29574 0 2686 7299
0 3064 1731 17637 0 1127 3751
59 247 209 8554 419 162 1907
55 518 1292 23660 599 1346 3715
0 299 361 8670 0 1958 5167
0 646 720 191 0 130 54
0 0 992 0 0 6083 15271
0 16280 3762 5633 0 263 116
524 25975 17002 14371 383 931 366
0 11400 1737 7432 0 802 445
A+BH
A+IHW
A+D
G
RNA Pol II HiChIP in HCT116 (Control vs Auxin)
J
IKAROS CTCF HiChIP (WT vs IKDN)
H
H3K27ac HiChIP in HaCaT (WT vs Stim)
K
IKAROS SMC1 HiChIP (WT vs IKDN)
I
H3K27ac HiChIP in melanoma cells (WT vs STAG2-KD)
L
IKAROS H3K27ac HiChIP (WT vs IKDN)
Union Significant FitHiChIP Loops
Loops in A+BH
Loops in A+IHW
Loops in A+D
Exclusive loops in A+IHW
Exclusive loops in A+D
Genomic Distance
0
0.5 Mb
1 Mb
1.5 Mb
2 Mb
3 Mb
4 Mb
Union Significant FitHiChIP Loops
DESeq2
edgeR exactTest
edgeR glmLRT
edgeR glmQLFTest

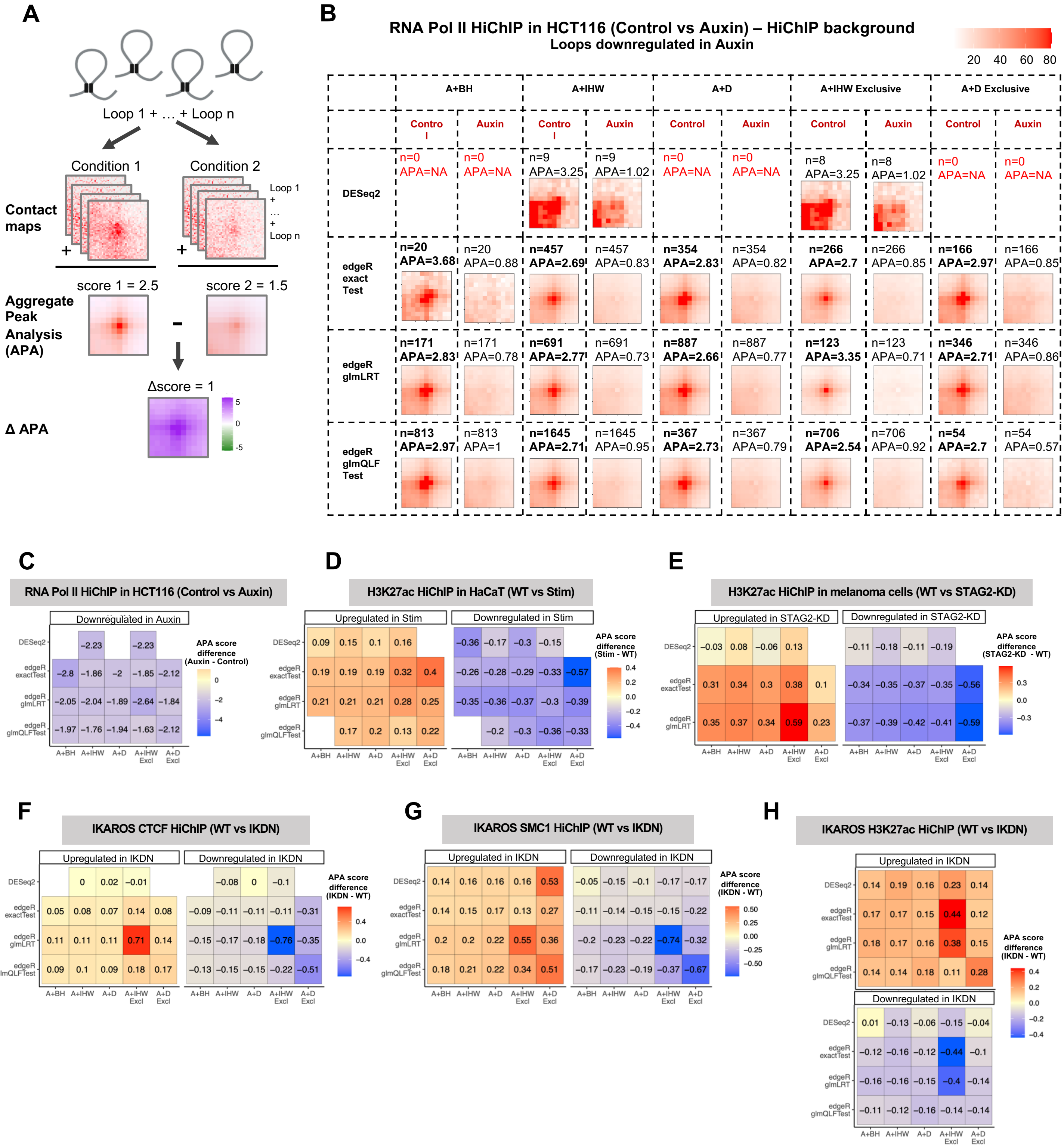

A
Loop 1 + … + Loop n
Condition 1
Condition 2
Contact maps
Loop 1
+
…
+
Loop n
score 1 = 2.5
score 2 = 1.5
Aggregate Peak Analysis (APA)
Δscore = 1
Δ APA
B
RNA Pol II HiChIP in HCT116 (Control vs Auxin) – HiChIP background
Loops downregulated in Auxin
20 40 60 80
A+BH
A+IHW
A+D
A+IHW Exclusive
A+D Exclusive
Control
Auxin
DESeq2
edgeR exact Test
edgeR glmLRT
edgeR glmQLF Test
n=0 APA=NA
n=9 APA=3.25
n=9 APA=1.02
n=8 APA=3.25
n=8 APA=1.02
n=20 APA=3.68
n=20 APA=0.88
n=457 APA=2.69
n=457 APA=0.83
n=354 APA=2.83
n=354 APA=0.82
n=266 APA=2.7
n=266 APA=0.85
n=166 APA=2.97
n=166 APA=0.85
n=171 APA=2.83
n=171 APA=0.78
n=691 APA=2.77
n=691 APA=0.73
n=887 APA=2.66
n=887 APA=0.77
n=123 APA=3.35
n=123 APA=0.71
n=346 APA=2.71
n=346 APA=0.86
n=813 APA=2.97
n=813 APA=1
n=1645 APA=2.71
n=1645 APA=0.95
n=367 APA=2.73
n=367 APA=0.79
n=706 APA=2.54
n=706 APA=0.92
n=54 APA=2.7
n=54 APA=0.57
C
RNA Pol II HiChIP in HCT116 (Control vs Auxin)
Downregulated in Auxin
APA score difference (Auxin - Control)
D
H3K27ac HiChIP in HaCaT (WT vs Stim)
Upregulated in Stim
Downregulated in Stim
APA score difference (Stim - WT)
E
H3K27ac HiChIP in melanoma cells (WT vs STAG2-KD)
Upregulated in STAG2-KD
Downregulated in STAG2-KD
APA score difference (STAG2-KD - WT)
F
IKAROS CTCF HiChIP (WT vs IKDN)
Upregulated in IKDN
Downregulated in IKDN
APA score difference (IKDN - WT)
G
IKAROS SMC1 HiChIP (WT vs IKDN)
H
IKAROS H3K27ac HiChIP (WT vs IKDN)
DESeq2
edgeR exactTest
edgeR glmLRT
edgeR glmQLFTest
A+BH
A+IHW
A+D
A+IHW Excl
A+D Excl

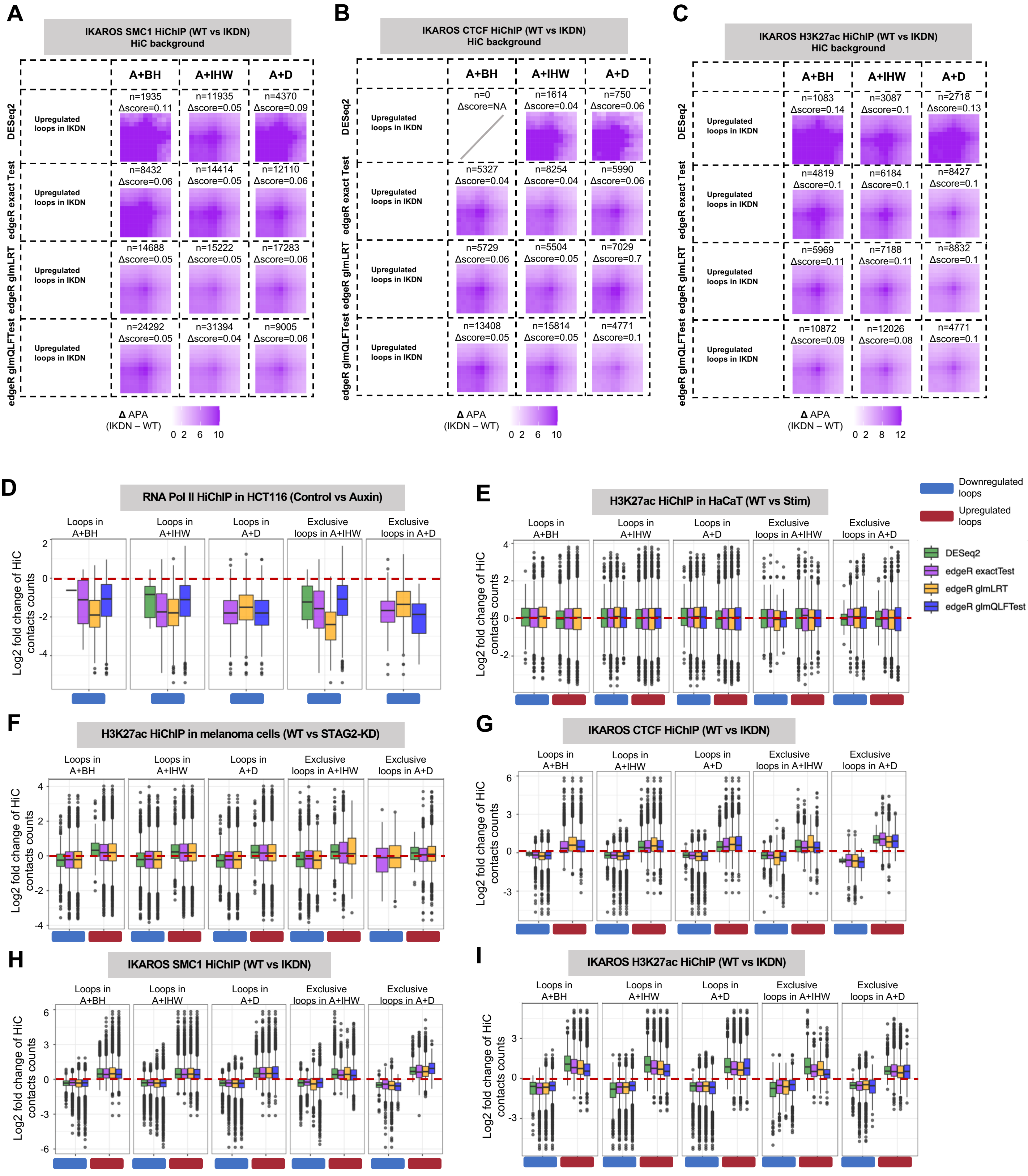
A
IKAROS SMC1 HiChIP (WT vs IKDN)
HiC background
A+BH
A+IHW
A+D
DESeq2
edgeR exact Test
edgeR glmLRT
edgeR glmQLFTest
Upregulated loops in IKDN
n=1935 Δscore=0.11
n=11935 Δscore=0.05
n=4370 Δscore=0.09
n=8432 Δscore=0.06
n=14414 Δscore=0.05
n=12110 Δscore=0.06
n=14688 Δscore=0.05
n=15222 Δscore=0.05
n=17283 Δscore=0.06
n=24292 Δscore=0.05
n=31394 Δscore=0.04
n=9005 Δscore=0.06
Δ APA (IKDN – WT)
0 2 6 10
B
IKAROS CTCF HiChIP (WT vs IKDN)
HiC background
n=0 Δscore=NA
n=1614 Δscore=0.04
n=750 Δscore=0.06
n=5327 Δscore=0.04
n=8254 Δscore=0.04
n=5990 Δscore=0.06
n=5729 Δscore=0.06
n=5504 Δscore=0.05
n=7029 Δscore=0.7
n=13408 Δscore=0.05
n=15814 Δscore=0.05
n=4771 Δscore=0.1
0 2 6 10
C
IKAROS H3K27ac HiChIP (WT vs IKDN)
HiC background
n=1083 Δscore=0.14
n=3087 Δscore=0.1
n=2718 Δscore=0.13
n=4819 Δscore=0.1
n=6184 Δscore=0.1
n=8427 Δscore=0.1
n=5969 Δscore=0.11
n=7188 Δscore=0.11
n=8832 Δscore=0.1
n=10872 Δscore=0.09
n=12026 Δscore=0.08
n=4771 Δscore=0.1
0 2 8 12
D
RNA Pol II HiChIP in HCT116 (Control vs Auxin)
E
H3K27ac HiChIP in HaCaT (WT vs Stim)
F
H3K27ac HiChIP in melanoma cells (WT vs STAG2-KD)
G
IKAROS CTCF HiChIP (WT vs IKDN)
H
IKAROS SMC1 HiChIP (WT vs IKDN)
I
IKAROS H3K27ac HiChIP (WT vs IKDN)
Loops in A+BH
Loops in A+IHW
Loops in A+D
Exclusive loops in A+IHW
Exclusive loops in A+D
Log2 fold change of HiC contacts counts
Downregulated loops
Upregulated loops
DESeq2
edgeR exactTest
edgeR glmLRT
edgeR glmQLFTest

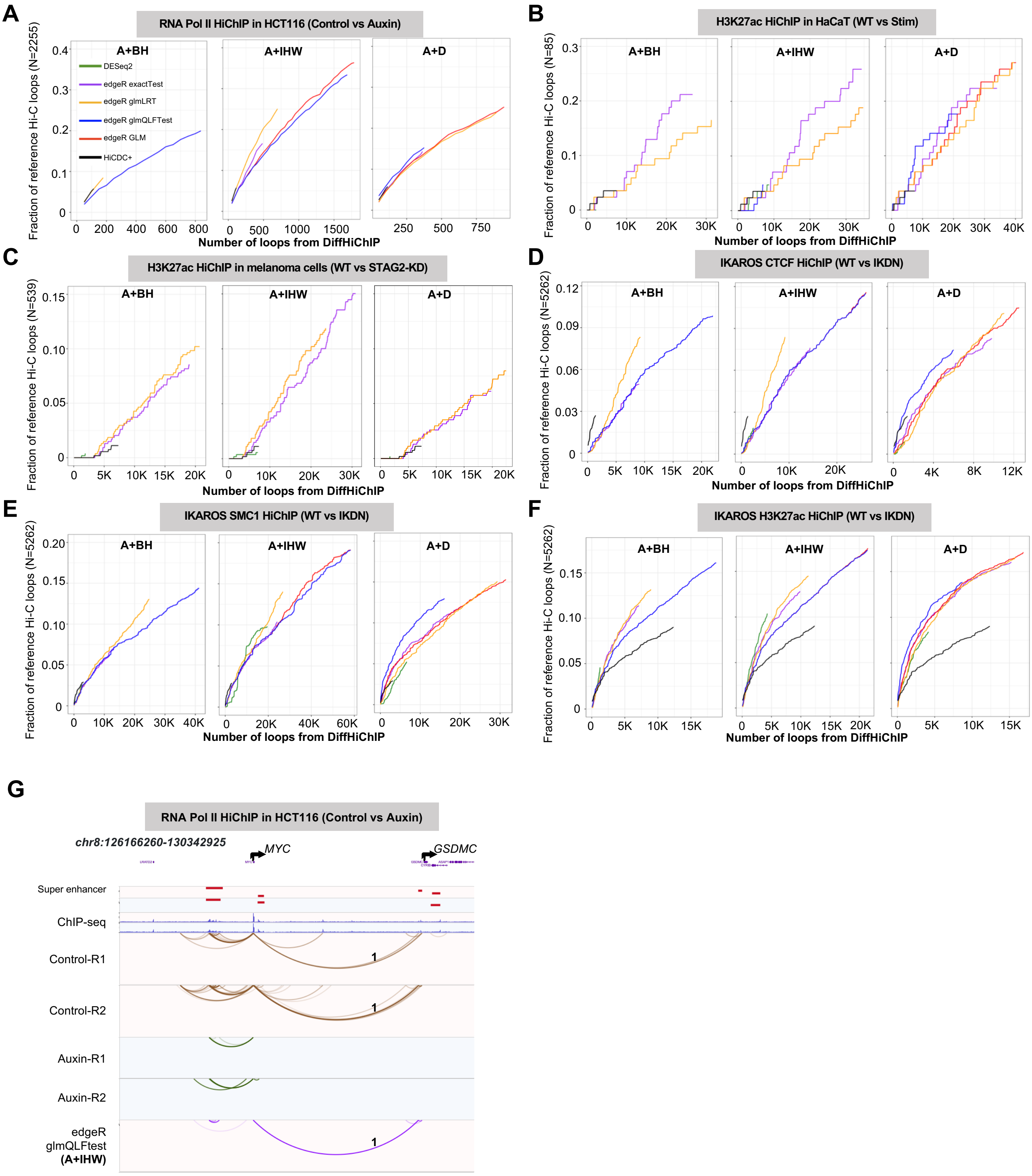

A
RNA Pol II HiChIP in HCT116 (Control vs Auxin)
A+BH
A+IHW
A+D
DESeq2
edgeR exactTest
edgeR glmLRT
edgeR glmQLFTest
edgeR GLM
HiCDC+
Fraction of reference Hi-C loops (N=2255)
Number of loops from DiffHiChIP
B
H3K27ac HiChIP in HaCaT (WT vs Stim)
Fraction of reference Hi-C loops (N=85)
C
H3K27ac HiChIP in melanoma cells (WT vs STAG2-KD)
Fraction of reference Hi-C loops (N=539)
D
IKAROS CTCF HiChIP (WT vs IKDN)
Fraction of reference Hi-C loops (N=5262)
E
IKAROS SMC1 HiChIP (WT vs IKDN)
F
IKAROS H3K27ac HiChIP (WT vs IKDN)
G
RNA Pol II HiChIP in HCT116 (Control vs Auxin)
chr8:126166260-130342925
MYC
GSDMC
Super enhancer
ChIP-seq
Control-R1
Control-R2
Auxin-R1
Auxin-R2
edgeR
glmQLFtest
(A+IHW)

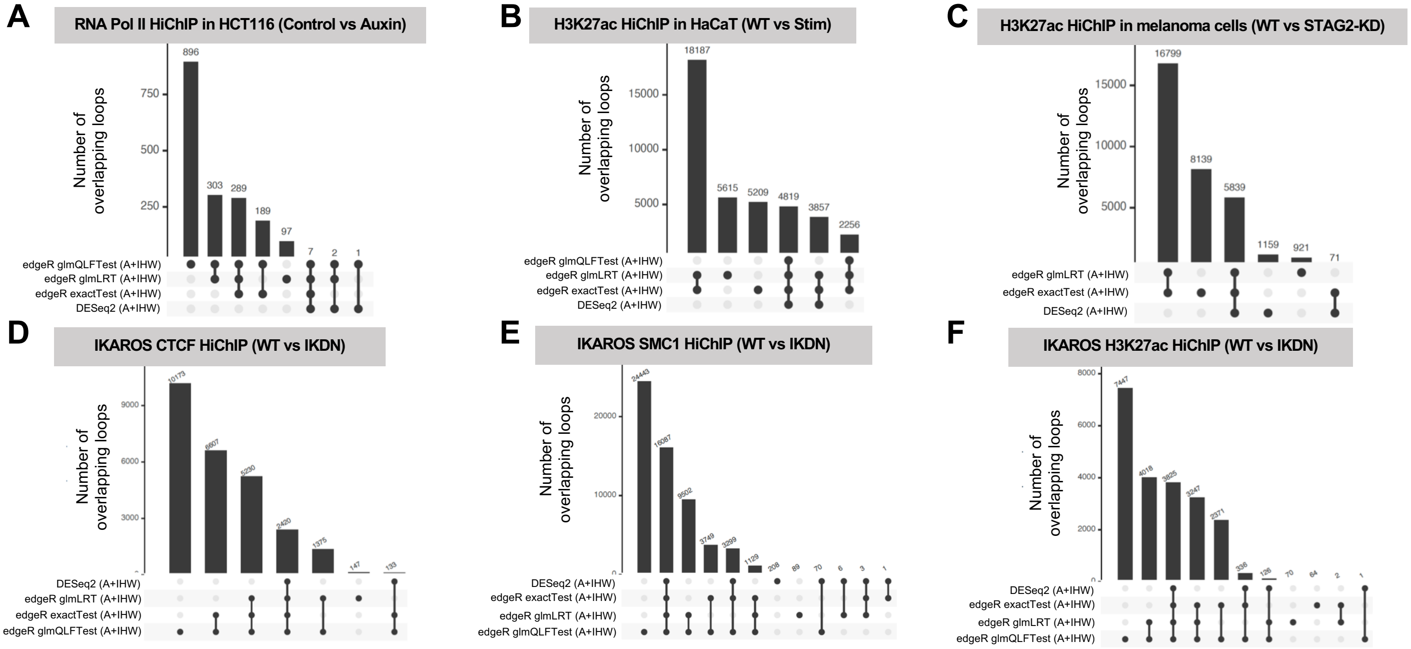
A
RNA Pol II HiChIP in HCT116 (Control vs Auxin)
Number of overlapping loops
edgeR glmQLFTest (A+IHW)
edgeR glmLRT (A+IHW)
edgeR exactTest (A+IHW)
DESeq2 (A+IHW)
896
303
289
189
97
7
2
1
B
H3K27ac HiChIP in HaCaT (WT vs Stim)
18187
5615
5209
4819
3857
2256
C
H3K27ac HiChIP in melanoma cells (WT vs STAG2-KD)
16799
8139
5839
1159
921
71
D
IKAROS CTCF HiChIP (WT vs IKDN)
E
IKAROS SMC1 HiChIP (WT vs IKDN)
F
IKAROS H3K27ac HiChIP (WT vs IKDN)

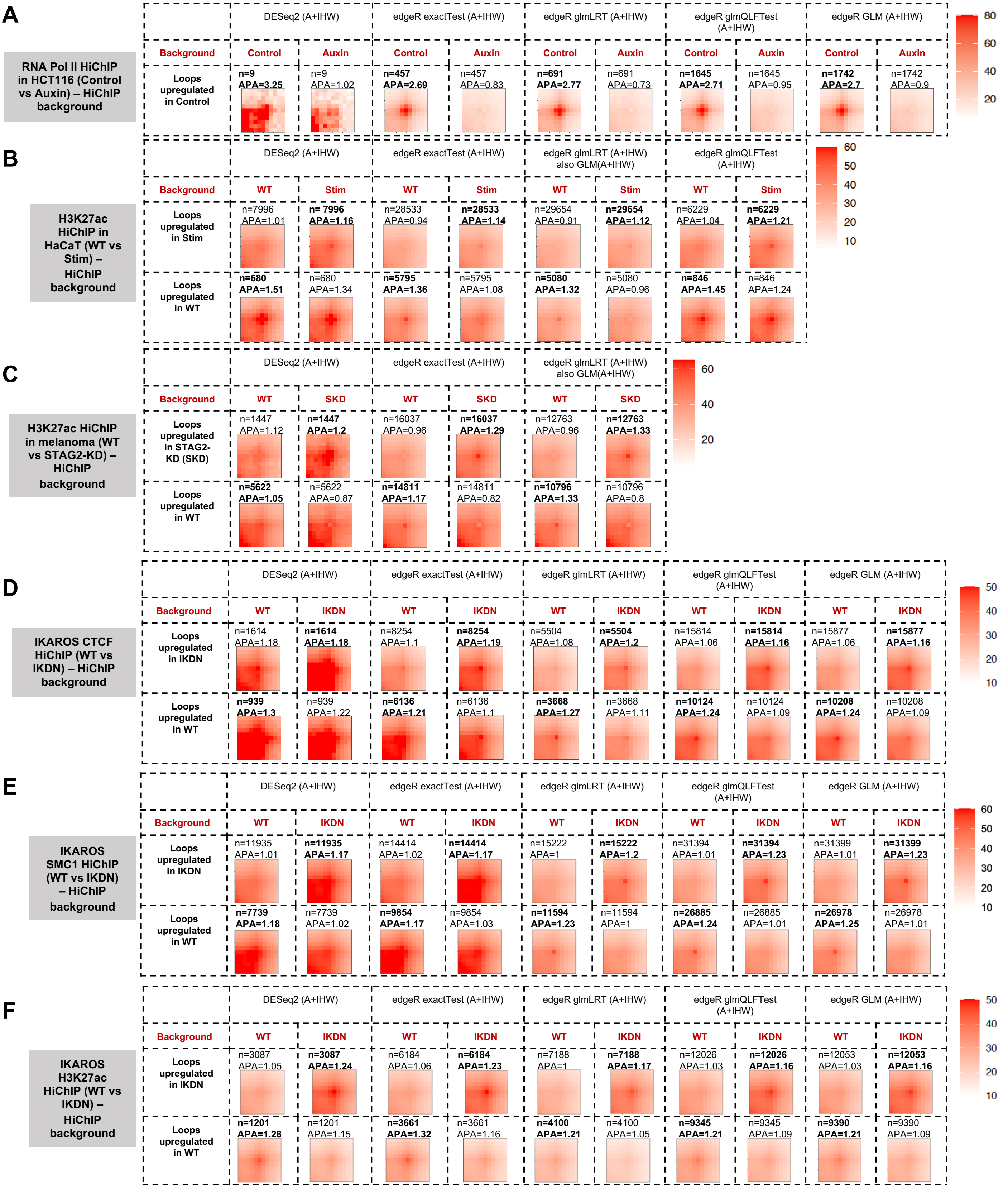
A
RNA Pol II HiChIP in HCT116 (Control vs Auxin) – HiChIP background
DESeq2 (A+IHW)
edgeR exactTest (A+IHW)
edgeR glmLRT (A+IHW)
edgeR glmQLFTest (A+IHW)
edgeR GLM (A+IHW)
Background
Control
Auxin
Loops upregulated in Control
n=9 APA=3.25
n=9 APA=1.02
n=457 APA=2.69
n=457 APA=0.83
n=691 APA=2.77
n=691 APA=0.73
n=1645 APA=2.71
n=1645 APA=0.95
n=1742 APA=2.7
n=1742 APA=0.9
80
60
40
20
B
H3K27ac HiChIP in HaCaT (WT vs Stim) – HiChIP background
DESeq2 (A+IHW)
edgeR exactTest (A+IHW)
edgeR glmLRT (A+IHW) also GLM(A+IHW)
edgeR glmQLFTest (A+IHW)
WT
Stim
Loops upregulated in Stim
n=7996 APA=1.01
n= 7996 APA=1.16
n=28533 APA=0.94
n=28533 APA=1.14
n=29654 APA=0.91
n=29654 APA=1.12
n=6229 APA=1.04
n=6229 APA=1.21
Loops upregulated in WT
n=680 APA=1.51
n=680 APA=1.34
n=5795 APA=1.36
n=5795 APA=1.08
n=5080 APA=1.32
n=5080 APA=0.96
n=846 APA=1.45
n=846 APA=1.24
60
50
40
30
20
10
C
H3K27ac HiChIP in melanoma (WT vs STAG2-KD) – HiChIP background
SKD
Loops upregulated in STAG2-KD (SKD)
n=1447 APA=1.12
n=1447 APA=1.2
n=16037 APA=0.96
n=16037 APA=1.29
n=12763 APA=0.96
n=12763 APA=1.33
n=5622 APA=1.05
n=5622 APA=0.87
n=14811 APA=1.17
n=14811 APA=0.82
n=10796 APA=1.33
n=10796 APA=0.8
60
40
20
D
IKAROS CTCF HiChIP (WT vs IKDN) – HiChIP background
IKDN
Loops upregulated in IKDN
n=1614 APA=1.18
n=1614 APA=1.18
n=8254 APA=1.1
n=8254 APA=1.19
n=5504 APA=1.08
n=5504 APA=1.2
n=15814 APA=1.06
n=15814 APA=1.16
n=15877 APA=1.06
n=15877 APA=1.16
n=939 APA=1.3
n=939 APA=1.22
n=6136 APA=1.21
n=6136 APA=1.1
n=3668 APA=1.27
n=3668 APA=1.11
n=10124 APA=1.24
n=10124 APA=1.09
n=10208 APA=1.24
n=10208 APA=1.09
50
40
30
20
10
E
IKAROS SMC1 HiChIP (WT vs IKDN) – HiChIP background
n=11935 APA=1.01
n=11935 APA=1.17
n=14414 APA=1.02
n=14414 APA=1.17
n=15222 APA=1
n=15222 APA=1.2
n=31394 APA=1.01
n=31394 APA=1.23
n=31399 APA=1.01
n=31399 APA=1.23
n=7739 APA=1.18
n=7739 APA=1.02
n=9854 APA=1.17
n=9854 APA=1.03
n=11594 APA=1.23
n=11594 APA=1
n=26885 APA=1.24
n=26885 APA=1.01
n=26978 APA=1.25
n=26978 APA=1.01
60
50
40
30
20
10
F
IKAROS H3K27ac HiChIP (WT vs IKDN) – HiChIP background
n=3087 APA=1.05
n=3087 APA=1.24
n=6184 APA=1.06
n=6184 APA=1.23
n=7188 APA=1
n=7188 APA=1.17
n=12026 APA=1.03
n=12026 APA=1.16
n=12053 APA=1.03
n=12053 APA=1.16
n=1201 APA=1.28
n=1201 APA=1.15
n=3661 APA=1.32
n=3661 APA=1.16
n=4100 APA=1.21
n=4100 APA=1.05
n=9345 APA=1.21
n=9345 APA=1.09
n=9390 APA=1.21
n=9390 APA=1.09
50
40
30
20
10

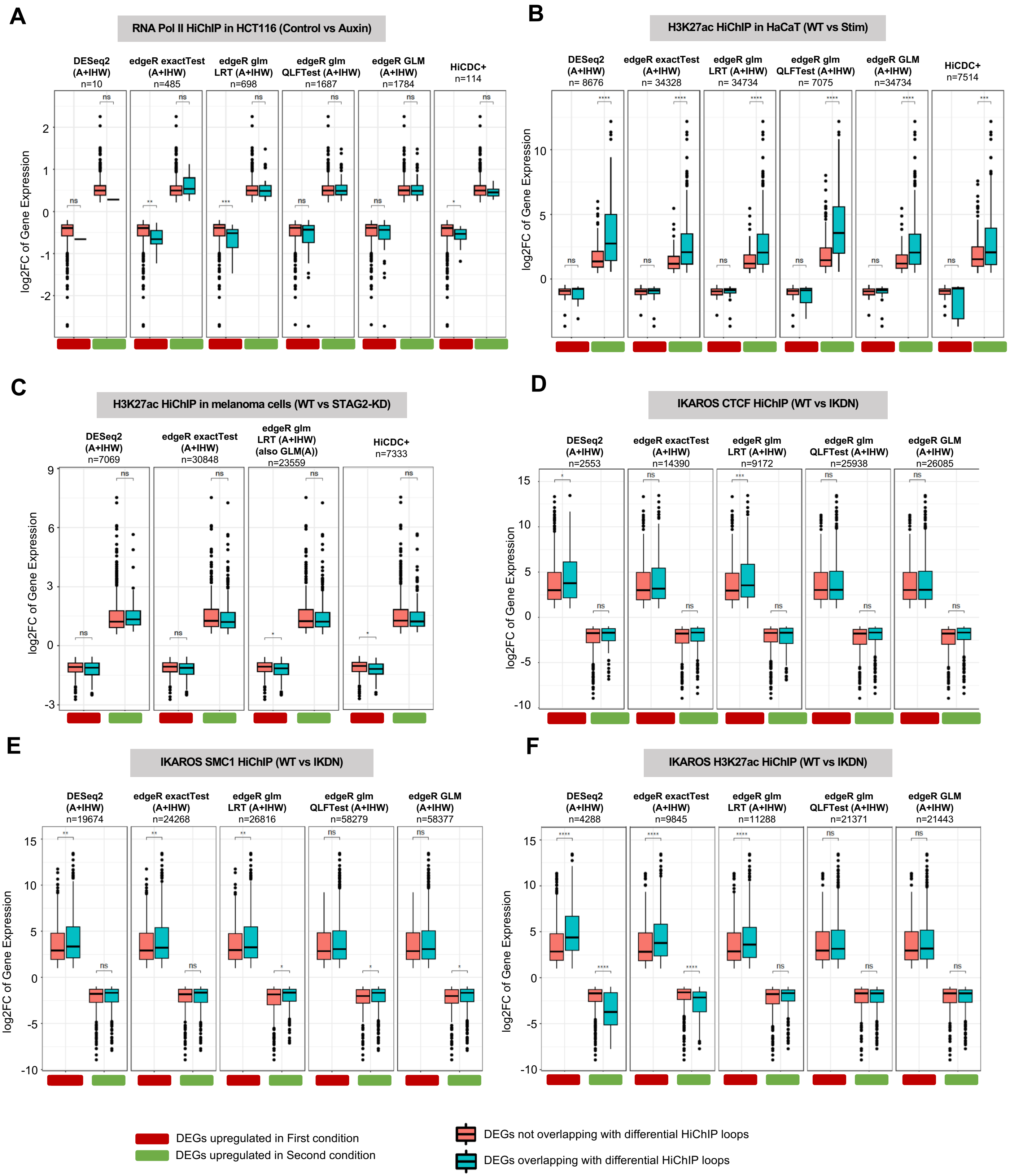
A
RNA Pol II HiChIP in HCT116 (Control vs Auxin)
DESeq2 (A+IHW) n=10
edgeR exactTest (A+IHW) n=485
edgeR glm LRT (A+IHW) n=698
edgeR glm QLFTest (A+IHW) n=1687
edgeR GLM (A+IHW) n=1784
HiCDC+ n=114
log2FC of Gene Expression
B
H3K27ac HiChIP in HaCaT (WT vs Stim)
DESeq2 (A+IHW) n= 8676
edgeR exactTest (A+IHW) n= 34328
edgeR glm LRT (A+IHW) n= 34734
edgeR glm QLFTest (A+IHW) n= 7075
edgeR GLM (A+IHW) n=34734
HiCDC+ n=7514
C
H3K27ac HiChIP in melanoma cells (WT vs STAG2-KD)
DESeq2 (A+IHW) n=7069
edgeR exactTest (A+IHW) n=30848
edgeR glm LRT (A+IHW) (also GLM(A)) n=23559
HiCDC+ n=7333
D
IKAROS CTCF HiChIP (WT vs IKDN)
DESeq2 (A+IHW) n=2553
edgeR exactTest (A+IHW) n=14390
edgeR glm LRT (A+IHW) n=9172
edgeR glm QLFTest (A+IHW) n=25938
edgeR GLM (A+IHW) n=26085
E
IKAROS SMC1 HiChIP (WT vs IKDN)
DESeq2 (A+IHW) n=19674
edgeR exactTest (A+IHW) n=24268
edgeR glm LRT (A+IHW) n=26816
edgeR glm QLFTest (A+IHW) n=58279
edgeR GLM (A+IHW) n=58377
F
IKAROS H3K27ac HiChIP (WT vs IKDN)
DESeq2 (A+IHW) n=4288
edgeR exactTest (A+IHW) n=9845
edgeR glm LRT (A+IHW) n=11288
edgeR glm QLFTest (A+IHW) n=21371
edgeR GLM (A+IHW) n=21443
DEGs upregulated in First condition
DEGs upregulated in Second condition
DEGs not overlapping with differential HiChIP loops
DEGs overlapping with differential HiChIP loops

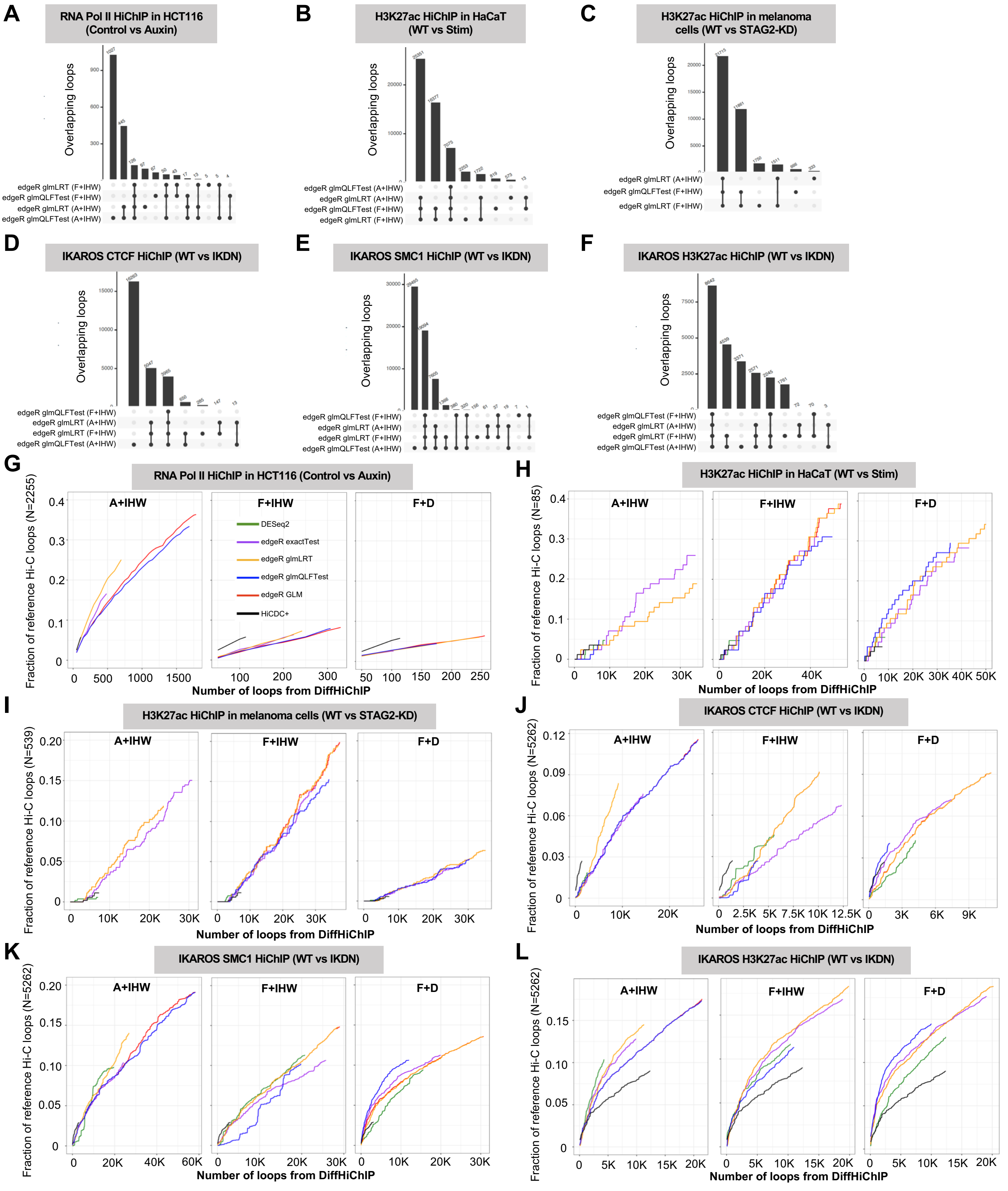

A
RNA Pol II HiChIP in HCT116 (Control vs Auxin)
B
H3K27ac HiChIP in HaCaT (WT vs Stim)
C
H3K27ac HiChIP in melanoma cells (WT vs STAG2-KD)
D
IKAROS CTCF HiChIP (WT vs IKDN)
E
IKAROS SMC1 HiChIP (WT vs IKDN)
F
IKAROS H3K27ac HiChIP (WT vs IKDN)
Overlapping loops
edgeR glmLRT (F+IHW)
edgeR glmQLFTest (F+IHW)
edgeR glmLRT (A+IHW)
edgeR glmQLFTest (A+IHW)
G
RNA Pol II HiChIP in HCT116 (Control vs Auxin)
H
H3K27ac HiChIP in HaCaT (WT vs Stim)
I
H3K27ac HiChIP in melanoma cells (WT vs STAG2-KD)
J
IKAROS CTCF HiChIP (WT vs IKDN)
K
IKAROS SMC1 HiChIP (WT vs IKDN)
L
IKAROS H3K27ac HiChIP (WT vs IKDN)
A+IHW
F+IHW
F+D
DESeq2
edgeR exactTest
edgeR glmLRT
edgeR glmQLFTest
edgeR GLM
HiCDC+
Fraction of reference Hi-C loops (N=2255)
Fraction of reference Hi-C loops (N=85)
Fraction of reference Hi-C loops (N=539)
Fraction of reference Hi-C loops (N=5262)
Number of loops from DiffHiChIP

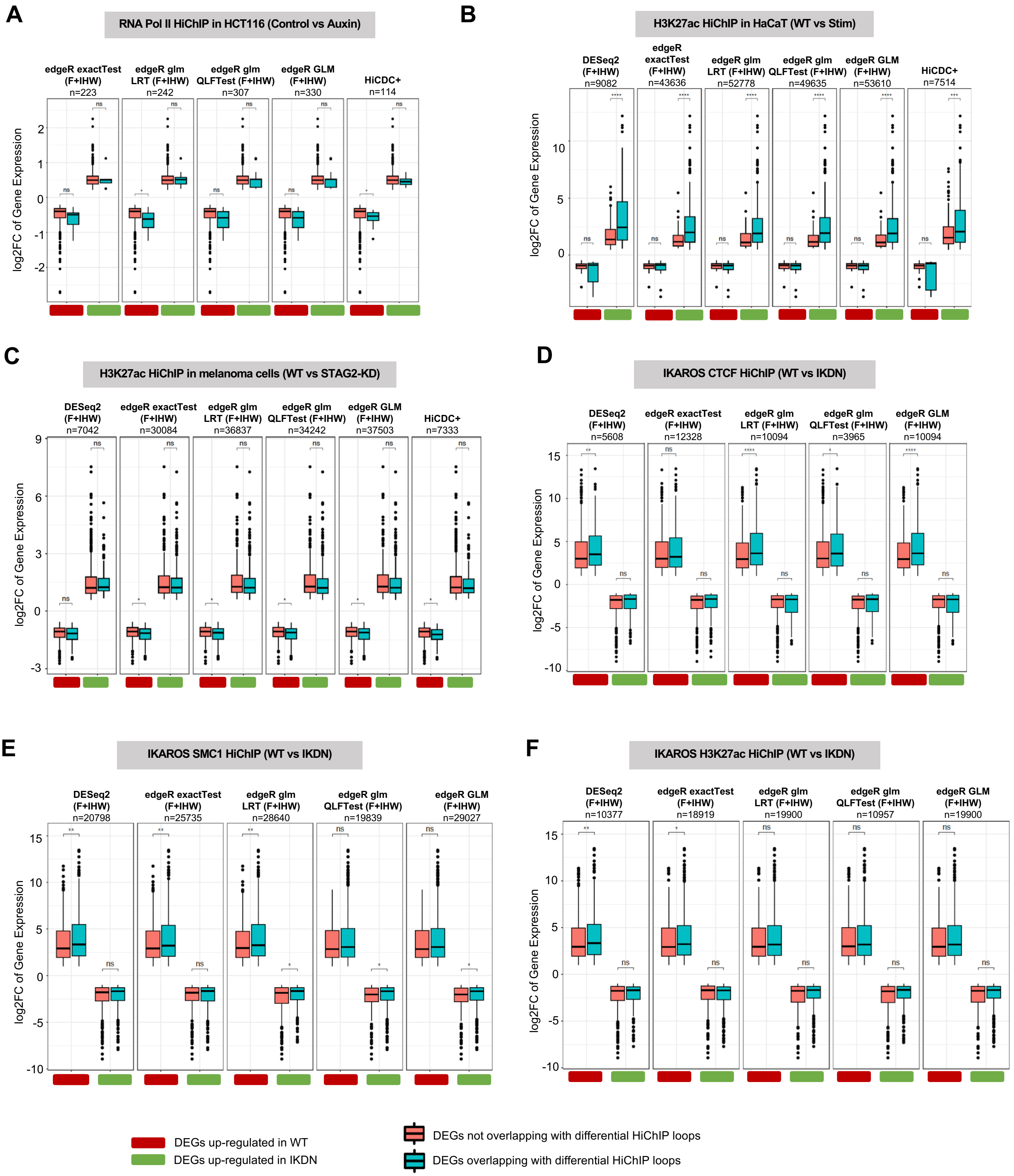

A
RNA Pol II HiChIP in HCT116 (Control vs Auxin)
edgeR exactTest (F+IHW) n=223
edgeR glm LRT (F+IHW) n=242
edgeR glm QLFTest (F+IHW) n=307
edgeR GLM (F+IHW) n=330
HiCDC+ n=114
log2FC of Gene Expression
B
H3K27ac HiChIP in HaCaT (WT vs Stim)
DESeq2 (F+IHW) n=9082
edgeR exactTest (F+IHW) n=43636
edgeR glm LRT (F+IHW) n=52778
edgeR glm QLFTest (F+IHW) n=49635
edgeR GLM (F+IHW) n=53610
HiCDC+ n=7514
log2FC of Gene Expression
C
H3K27ac HiChIP in melanoma cells (WT vs STAG2-KD)
DESeq2 (F+IHW) n=7042
edgeR exactTest (F+IHW) n=30084
edgeR glm LRT (F+IHW) n=36837
edgeR glm QLFTest (F+IHW) n=34242
edgeR GLM (F+IHW) n=37503
HiCDC+ n=7333
log2FC of Gene Expression
D
IKAROS CTCF HiChIP (WT vs IKDN)
DESeq2 (F+IHW) n=5608
edgeR exactTest (F+IHW) n=12328
edgeR glm LRT (F+IHW) n=10094
edgeR glm QLFTest (F+IHW) n=3965
edgeR GLM (F+IHW) n=10094
log2FC of Gene Expression
E
IKAROS SMC1 HiChIP (WT vs IKDN)
DESeq2 (F+IHW) n=20798
edgeR exactTest (F+IHW) n=25735
edgeR glm LRT (F+IHW) n=28640
edgeR glm QLFTest (F+IHW) n=19839
edgeR GLM (F+IHW) n=29027
log2FC of Gene Expression
F
IKAROS H3K27ac HiChIP (WT vs IKDN)
DESeq2 (F+IHW) n=10377
edgeR exactTest (F+IHW) n=18919
edgeR glm LRT (F+IHW) n=19900
edgeR glm QLFTest (F+IHW) n=10957
edgeR GLM (F+IHW) n=19900
log2FC of Gene Expression
DEGs up-regulated in WT
DEGs up-regulated in IKDN
DEGs not overlapping with differential HiChIP loops
DEGs overlapping with differential HiChIP loops

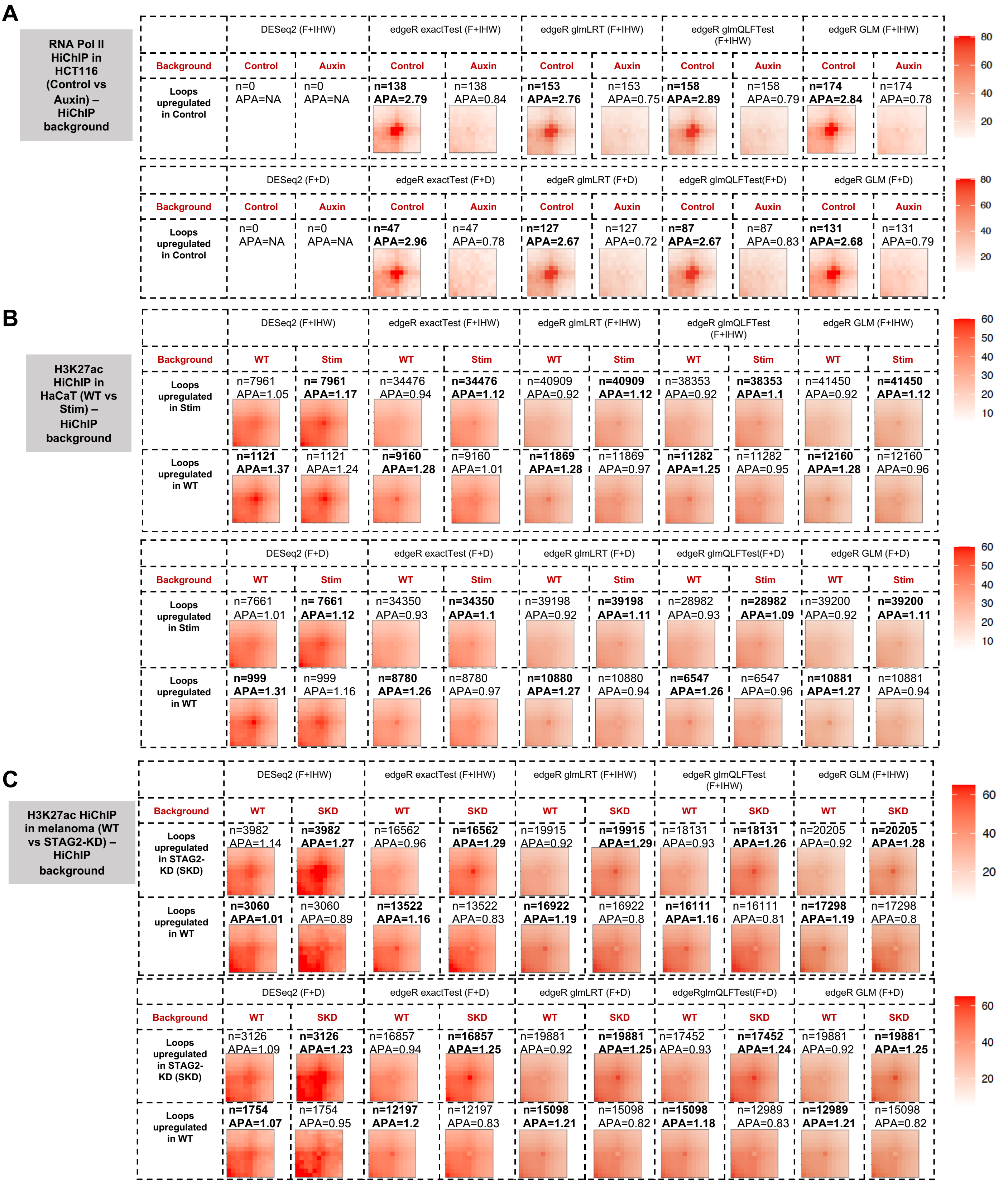

A
RNA Pol II HiChIP in HCT116 (Control vs Auxin) – HiChIP background
DESeq2 (F+IHW)
edgeR exactTest (F+IHW)
edgeR glmLRT (F+IHW)
edgeR glmQLFTest (F+IHW)
edgeR GLM (F+IHW)
Background
Control
Auxin
Loops upregulated in Control
n=0 APA=NA
n=0 APA=NA
n=138 APA=2.79
n=138 APA=0.84
n=153 APA=2.76
n=153 APA=0.75
n=158 APA=2.89
n=158 APA=0.79
n=174 APA=2.84
n=174 APA=0.78
80
60
40
20
DESeq2 (F+D)
edgeR exactTest (F+D)
edgeR glmLRT (F+D)
edgeR glmQLFTest(F+D)
edgeR GLM (F+D)
Background
Control
Auxin
Loops upregulated in Control
n=0 APA=NA
n=0 APA=NA
n=47 APA=2.96
n=47 APA=0.78
n=127 APA=2.67
n=127 APA=0.72
n=87 APA=2.67
n=87 APA=0.83
n=131 APA=2.68
n=131 APA=0.79
80
60
40
20
B
H3K27ac HiChIP in HaCaT (WT vs Stim) – HiChIP background
DESeq2 (F+IHW)
edgeR exactTest (F+IHW)
edgeR glmLRT (F+IHW)
edgeR glmQLFTest (F+IHW)
edgeR GLM (F+IHW)
Background
WT
Stim
Loops upregulated in Stim
n=7961 APA=1.05
n= 7961 APA=1.17
n=34476 APA=0.94
n=34476 APA=1.12
n=40909 APA=0.92
n=40909 APA=1.12
n=38353 APA=0.92
n=38353 APA=1.1
n=41450 APA=0.92
n=41450 APA=1.12
Loops upregulated in WT
n=1121 APA=1.37
n=1121 APA=1.24
n=9160 APA=1.28
n=9160 APA=1.01
n=11869 APA=1.28
n=11869 APA=0.97
n=11282 APA=1.25
n=11282 APA=0.95
n=12160 APA=1.28
n=12160 APA=0.96
60
50
40
30
20
10
DESeq2 (F+D)
edgeR exactTest (F+D)
edgeR glmLRT (F+D)
edgeR glmQLFTest(F+D)
edgeR GLM (F+D)
Background
WT
Stim
Loops upregulated in Stim
n=7661 APA=1.01
n= 7661 APA=1.12
n=34350 APA=0.93
n=34350 APA=1.1
n=39198 APA=0.92
n=39198 APA=1.11
n=28982 APA=0.93
n=28982 APA=1.09
n=39200 APA=0.92
n=39200 APA=1.11
Loops upregulated in WT
n=999 APA=1.31
n=999 APA=1.16
n=8780 APA=1.26
n=8780 APA=0.97
n=10880 APA=1.27
n=10880 APA=0.94
n=6547 APA=1.26
n=6547 APA=0.96
n=10881 APA=1.27
n=10881 APA=0.94
60
50
40
30
20
10
C
H3K27ac HiChIP in melanoma (WT vs STAG2-KD) – HiChIP background
DESeq2 (F+IHW)
edgeR exactTest (F+IHW)
edgeR glmLRT (F+IHW)
edgeR glmQLFTest (F+IHW)
edgeR GLM (F+IHW)
Background
WT
SKD
Loops upregulated in STAG2-KD (SKD)
n=3982 APA=1.14
n=3982 APA=1.27
n=16562 APA=0.96
n=16562 APA=1.29
n=19915 APA=0.92
n=19915 APA=1.29
n=18131 APA=0.93
n=18131 APA=1.26
n=20205 APA=0.92
n=20205 APA=1.28
Loops upregulated in WT
n=3060 APA=1.01
n=3060 APA=0.89
n=13522 APA=1.16
n=13522 APA=0.83
n=16922 APA=1.19
n=16922 APA=0.8
n=16111 APA=1.16
n=16111 APA=0.81
n=17298 APA=1.19
n=17298 APA=0.8
60
40
20
DESeq2 (F+D)
edgeR exactTest (F+D)
edgeR glmLRT (F+D)
edgeRglmQLFTest(F+D)
edgeR GLM (F+D)
Background
WT
SKD
Loops upregulated in STAG2-KD (SKD)
n=3126 APA=1.09
n=3126 APA=1.23
n=16857 APA=0.94
n=16857 APA=1.25
n=19881 APA=0.92
n=19881 APA=1.25
n=17452 APA=0.93
n=17452 APA=1.24
n=19881 APA=0.92
n=19881 APA=1.25
Loops upregulated in WT
n=1754 APA=1.07
n=1754 APA=0.95
n=12197 APA=1.2
n=12197 APA=0.83
n=15098 APA=1.21
n=15098 APA=0.82
n=15098 APA=1.18
n=12989 APA=0.83
n=12989 APA=1.21
n=15098 APA=0.82
60
40
20

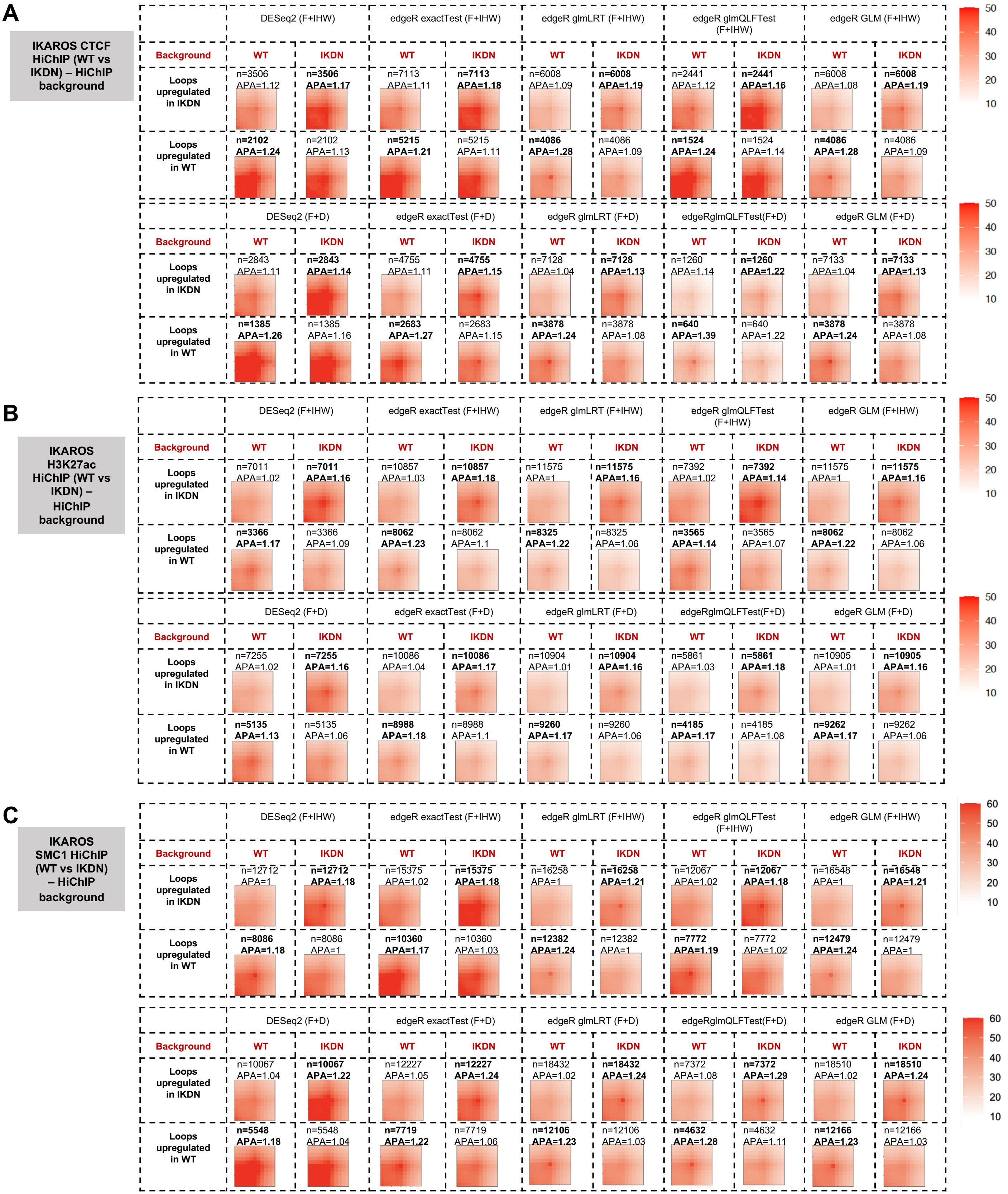
A
IKAROS CTCF HiChIP (WT vs IKDN) – HiChIP background
DESeq2 (F+IHW)
edgeR exactTest (F+IHW)
edgeR glmLRT (F+IHW)
edgeR glmQLFTest (F+IHW)
edgeR GLM (F+IHW)
Background
WT
IKDN
Loops upregulated in IKDN
n=3506 APA=1.12
n=3506 APA=1.17
n=7113 APA=1.11
n=7113 APA=1.18
n=6008 APA=1.09
n=6008 APA=1.19
n=2441 APA=1.12
n=2441 APA=1.16
n=6008 APA=1.08
n=6008 APA=1.19
Loops upregulated in WT
n=2102 APA=1.24
n=2102 APA=1.13
n=5215 APA=1.21
n=5215 APA=1.11
n=4086 APA=1.28
n=4086 APA=1.09
n=1524 APA=1.24
n=1524 APA=1.14
n=4086 APA=1.28
n=4086 APA=1.09
50
40
30
20
10
DESeq2 (F+D)
edgeR exactTest (F+D)
edgeR glmLRT (F+D)
edgeRglmQLFTest(F+D)
edgeR GLM (F+D)
Background
WT
IKDN
Loops upregulated in IKDN
n=2843 APA=1.11
n=2843 APA=1.14
n=4755 APA=1.11
n=4755 APA=1.15
n=7128 APA=1.04
n=7128 APA=1.13
n=1260 APA=1.14
n=1260 APA=1.22
n=7133 APA=1.04
n=7133 APA=1.13
Loops upregulated in WT
n=1385 APA=1.26
n=1385 APA=1.16
n=2683 APA=1.27
n=2683 APA=1.15
n=3878 APA=1.24
n=3878 APA=1.08
n=640 APA=1.39
n=640 APA=1.22
n=3878 APA=1.24
n=3878 APA=1.08
50
40
30
20
10
B
IKAROS H3K27ac HiChIP (WT vs IKDN) – HiChIP background
DESeq2 (F+IHW)
edgeR exactTest (F+IHW)
edgeR glmLRT (F+IHW)
edgeR glmQLFTest (F+IHW)
edgeR GLM (F+IHW)
Background
WT
IKDN
Loops upregulated in IKDN
n=7011 APA=1.02
n=7011 APA=1.16
n=10857 APA=1.03
n=10857 APA=1.18
n=11575 APA=1
n=11575 APA=1.16
n=7392 APA=1.02
n=7392 APA=1.14
n=11575 APA=1
n=11575 APA=1.16
Loops upregulated in WT
n=3366 APA=1.17
n=3366 APA=1.09
n=8062 APA=1.23
n=8062 APA=1.1
n=8325 APA=1.22
n=8325 APA=1.06
n=3565 APA=1.14
n=3565 APA=1.07
n=8062 APA=1.22
n=8062 APA=1.06
50
40
30
20
10
DESeq2 (F+D)
edgeR exactTest (F+D)
edgeR glmLRT (F+D)
edgeRglmQLFTest(F+D)
edgeR GLM (F+D)
Background
WT
IKDN
Loops upregulated in IKDN
n=7255 APA=1.02
n=7255 APA=1.16
n=10086 APA=1.04
n=10086 APA=1.17
n=10904 APA=1.01
n=10904 APA=1.16
n=5861 APA=1.03
n=5861 APA=1.18
n=10905 APA=1.01
n=10905 APA=1.16
Loops upregulated in WT
n=5135 APA=1.13
n=5135 APA=1.06
n=8988 APA=1.18
n=8988 APA=1.1
n=9260 APA=1.17
n=9260 APA=1.06
n=4185 APA=1.17
n=4185 APA=1.08
n=9262 APA=1.17
n=9262 APA=1.06
50
40
30
20
10
C
IKAROS SMC1 HiChIP (WT vs IKDN) – HiChIP background
DESeq2 (F+IHW)
edgeR exactTest (F+IHW)
edgeR glmLRT (F+IHW)
edgeR glmQLFTest (F+IHW)
edgeR GLM (F+IHW)
Background
WT
IKDN
Loops upregulated in IKDN
n=12712 APA=1
n=12712 APA=1.18
n=15375 APA=1.02
n=15375 APA=1.18
n=16258 APA=1
n=16258 APA=1.21
n=12067 APA=1.02
n=12067 APA=1.18
n=16548 APA=1
n=16548 APA=1.21
Loops upregulated in WT
n=8086 APA=1.18
n=8086 APA=1
n=10360 APA=1.17
n=10360 APA=1.03
n=12382 APA=1.24
n=12382 APA=1
n=7772 APA=1.19
n=7772 APA=1.02
n=12479 APA=1.24
n=12479 APA=1
60
50
40
30
20
10
DESeq2 (F+D)
edgeR exactTest (F+D)
edgeR glmLRT (F+D)
edgeRglmQLFTest(F+D)
edgeR GLM (F+D)
Background
WT
IKDN
Loops upregulated in IKDN
n=10067 APA=1.04
n=10067 APA=1.22
n=12227 APA=1.05
n=12227 APA=1.24
n=18432 APA=1.02
n=18432 APA=1.24
n=7372 APA=1.08
n=7372 APA=1.29
n=18510 APA=1.02
n=18510 APA=1.24
Loops upregulated in WT
n=5548 APA=1.18
n=5548 APA=1.04
n=7719 APA=1.22
n=7719 APA=1.06
n=12106 APA=1.23
n=12106 APA=1.03
n=4632 APA=1.28
n=4632 APA=1.11
n=12166 APA=1.23
n=12166 APA=1.03
60
50
40
30
20
10
