## Supplementary Figure Legends for "DiffHiChIP: Identifying differential chromatin contacts from HiChIP data"

**Supplementary Figure 1: (A-F)** Overlap of differential loops between BH (A+BH) corrected FDR, IHW corrected FDR (A+IHW) and distance stratification (A+D) for various DESeq2 and edgeR settings and HiChIP datasets: HCT116 dataset for Control vs Auxin treated conditions **(A)**, HaCaT dataset for WT vs stimulated conditions **(B)**, Melanoma dataset for WT vs STAG2 knockdown (KD) conditions **(C)**, IKAROS CTCF dataset for WT vs IKDN conditions **(D)**, IKAROS SMC1 dataset for WT vs IKDN conditions **(E)** and IKAROS H3K27ac dataset for WT vs IKDN conditions **(F)**. Here, DiffHiChIP is executed in the complete background (A) setting**. (G-L)** Genomic distance of differential loops for different DiffHiChIP settings and for various HiChIP datasets: HCT116 dataset for Control vs Auxin treated conditions **(G)**, HaCaT dataset for WT vs stimulated conditions **(H)**, Melanoma dataset for WT vs STAG2 knockdown (KD) conditions **(I)**, IKAROS CTCF dataset for WT vs IKDN conditions **(J)**, IKAROS SMC1 dataset for WT vs IKDN conditions **(K)** and IKAROS H3K27ac dataset for WT vs IKDN conditions **(L)**.

**Supplementary Figure 2:** **(A)** Schematic illustrates the aggregate peak analysis (APA) and differential APA (Δ APA — APA matrix of one condition subtracted from the other) from a starting set of differential loops. The symbol “n” denotates the number of differential loops to be aggregated. Differential APA scores (Δscore) represent the difference in APA scores between conditions. **(B)** APA for downregulated loops in Auxin obtained by FDR with BH (A+BH) and IHW (A+IHW) corrections and distance stratification (A+D), and also the loops exclusively detected A+IHW and A+D, corresponding to the HCT116 HiChIP datasets between Control and Auxin conditions. Here HiChIP contacts for the same conditions are used as the backgrounds. Values in bold denote expectation of higher APAs among the two conditions (i.e., upregulated loops in that conditions). **(C-H)** Heatmap with APA score differences between the two conditions. Higher magnitude of differential APA scores indicates higher enrichment of loops in the respective conditions. Differential loops were obtained by FDR with BH (A+BH) and IHW (A+IHW) corrections and distance stratification (A+D), and also the loops exclusively detected for A+IHW (A+IHW Excl) and A+D IHW (A+D Excl), corresponding to the various HiChIP datasets.

**Supplementary Figure 3:** **(A-C)** Differential APA plots (elementwise subtraction of the aggregate matrix for IKDN from that of WT) for IKAROS SMC1 **(A)**, CTCF **(B)** and H3K27ac **(C)** HiChIP datasets for various distance stratification settings, using Hi-C data as background. Differential APA scores (Δscore) between conditions represent the difference in APA scores between the IKDN and WT backgrounds. Higher magnitude of differential APA scores indicates higher enrichment of loops in the respective conditions **(D-I)** Log2 fold change in Hi-C contact counts for upregulated and downregulated loops for different DiffHiChIP settings and for various HiChIP datasets: HCT116 dataset for Control vs Auxin treated conditions **(D)**, HaCaT dataset for WT vs stimulated conditions **(E)**, Melanoma dataset for WT vs STAG2 knockdown (KD) conditions **(F)**, IKAROS CTCF dataset for WT vs IKDN conditions **(G)**, IKAROS SMC1 dataset for WT vs IKDN conditions **(H)** and IKAROS H3K27ac dataset for WT vs IKDN conditions **(I)**. Here, DiffHiChIP is executed in the complete background (A) setting.

**Supplementary Figure 4: (A-F)** Recovery of differential Hi-C loops (computed using FitHiC2 and applying a fold change condition) by different settings of DiffHiChIP and the reference method HiCDC+ for various HiChIP datasets. DiffHiChIP is executed with the complete background (A) setting. The symbol “N” indicates the number of reference Hi-C loops. The HiChIP datasets employed are: HCT116 dataset for Control vs Auxin treated conditions **(A)**, HaCaT dataset for WT vs stimulated conditions **(B)**, Melanoma dataset for WT vs STAG2 knockdown (KD) conditions **(C)**, IKAROS CTCF dataset for WT vs IKDN conditions **(D)**, IKAROS SMC1 dataset for WT vs IKDN conditions **(E)** and IKAROS H3K27ac dataset for WT vs IKDN conditions **(F)**. **(G)** Differential loops lost upon CTCF depletion (Auxin) that linked the gene *MYC* and a ~1.9Mb downstream super enhancer near the gene *GSDMC* (marked 1). This loop was detected as differential by only edgeR glmQLFTest for the A+IHW setting.

**Supplementary Figure 5:** **(A-F)** Overlap of differential loops between DESeq2 and various edgeR settings with respect to the complete background (A) with IHW-corrected FDR (A+IHW), and for different HiChIP datasets: HCT116 dataset for Control vs Auxin treated conditions **(A)**, HaCaT dataset for WT vs stimulated conditions **(B)**, Melanoma dataset for WT vs STAG2 knockdown (KD) conditions **(C)**, IKAROS CTCF dataset for WT vs IKDN conditions **(D)**, IKAROS SMC1 dataset for WT vs IKDN conditions **(E)** and IKAROS H3K27ac dataset for WT vs IKDN conditions **(F)**.

**Supplementary Figure 6: (A-F)** APA plots for various HiChIP datasets between respective conditions, for DESeq2 and different edgeR settings. All these models use complete background (A) and IHW-corrected FDR (A+IHW). Values in bold denote expectation of higher APAs among the two conditions (i.e., upregulated loops in that conditions). The HiChIP datasets employed are: HCT116 dataset for Control vs Auxin treated conditions **(A)**, HaCaT dataset for WT vs stimulated conditions **(B)**, Melanoma dataset for WT vs STAG2 knockdown (KD) conditions **(C)**, IKAROS CTCF dataset for WT vs IKDN conditions **(D)**, IKAROS SMC1 dataset for WT vs IKDN conditions **(E)**, IKAROS H3K27ac dataset for WT vs IKDN conditions **(F)**. The symbol “n” indicates the number of differential loops.

**Supplementary Figure 7:** **(A-F)** Enrichment of magnitude of gene expression change (log2 fold change) for differential genes segregated with respect to their overlap with differential loops from different settings of DiffHiChIP (DESeq2 or edgeR), complete background (A) with IHW-corrected FDR (A+IHW), and for various HiChIP datasets: HCT116 dataset for Control vs Auxin treated conditions **(A)**, HaCaT dataset for WT vs stimulated conditions **(B)**, Melanoma dataset for WT vs STAG2 knockdown (KD) conditions **(C)**, IKAROS CTCF dataset for WT vs IKDN conditions **(D)**, IKAROS SMC1 dataset for WT vs IKDN conditions **(E)** and IKAROS H3K27ac dataset for WT vs IKDN conditions **(F)**. Enrichment is computed separately for genes upregulated in either condition. The symbol “n” indicates the number of differential loops. Significance was calculated using a Wilcoxon test (two-sided). *P<=0.05; **P<=0.01; ***P<=0.001; ****P<=0.0001; ns, not significant.

**Supplementary Figure 8:** **(A-F)** Overlap of differential loops for various edgeR GLM settings between the complete (A+IHW) and filtered (F+IHW) background settings of DiffHiChIP (with IHW), for various HiChIP datasets: HCT116 dataset for Control vs Auxin treated conditions **(A)**, HaCaT dataset for WT vs stimulated conditions **(B)**, Melanoma dataset for WT vs STAG2 knockdown (KD) conditions **(C)**, IKAROS CTCF dataset for WT vs IKDN conditions **(D)**, IKAROS SMC1 dataset for WT vs IKDN conditions **(E)** and IKAROS H3K27ac dataset for WT vs IKDN conditions **(F)**. **(G-L)** Recovery of differential Hi-C loops (computed using FitHiC2) by different settings of DiffHiChIP and the reference method HiCDC+ for various HiChIP datasets. DiffHiChIP is executed with the complete background (A) and filtered (F) setting, specifically A+IHW, F+IHW and F+D. The symbol “N” indicates the number of reference Hi-C loops. The HiChIP datasets employed are: HCT116 dataset for Control vs Auxin treated conditions **(G)**, HaCaT dataset for WT vs stimulated conditions **(H)**, Melanoma dataset for WT vs STAG2 knockdown (KD) conditions (I), IKAROS CTCF dataset for WT vs IKDN conditions **(J)**, IKAROS SMC1 dataset for WT vs IKDN conditions **(K)** and IKAROS H3K27ac dataset for WT vs IKDN conditions **(L)**.

**Supplementary Figure 9:** **(A-F)** Enrichment of magnitude of gene expression change (log2 fold change) for differential genes segregated with respect to their overlap with differential loops from different settings of DiffHiChIP (DESeq2 or edgeR) and with respect to the filtered background (F+IHW) for various HiChIP datasets: HCT116 dataset for Control vs Auxin treated conditions **(A)**, HaCaT dataset for WT vs stimulated conditions **(B)**, Melanoma dataset for WT vs STAG2 knockdown (KD) conditions **(C)**, IKAROS CTCF dataset for WT vs IKDN conditions **(D)**, IKAROS SMC1 dataset for WT vs IKDN conditions **(E)** and IKAROS H3K27ac dataset for WT vs IKDN conditions **(F)**. Enrichment is computed separately for genes upregulated in either condition. The The symbol “n” indicates the number of differential loops. Significance was calculated using a Wilcoxon test (two-sided). *P<=0.05; **P<=0.01; ***P<=0.001; ****P<=0.0001; ns, not significant.

**Supplementary Figure 10: (A-C)** APA plots for DiffHiChIP loops detected using filtered background and IHW corrected p-values (F+IHW) and for different DESeq2 and edgeR settings (exactTest, glmLRT, glmQLFTest and GLM) for different HiChIP datasets: HCT116 **(A)**, HaCaT **(B)** and Melanoma **(C)**. Values in bold denote expectation of higher APAs among the two conditions (i.e., loops upregulated in that condition).

**Supplementary Figure 11: (A-C)** Similar to Supplementary Figure 10 for various IKAROS HiChIP datasets: CTCF **(A)**, H3K27ac **(B)** and SMC1 **(C)**.
